# Ecological Diversification into Broad Thermal Niches Revealed by Protein Resurrection and Proteome-Wide Ancestral Reconstruction During the Rapid Radiation of Alvinellid Worms

**DOI:** 10.64898/2026.07.28.741189

**Authors:** Pierre-Guillaume Brun, Anne-Sophie Le Port, Marion Ballenghien, Sébastien Brûlé, Magali Aumont-Niçaise, Lionel Cladière, Didier Jollivet, Jean Mary

## Abstract

In the deep Pacific and Indian oceans, Alvinellid worms diversified about 100 million years ago to colonize a variety of hydrothermal vent environments. It has been suggested that the last common ancestor of this family was a thermophilic species. However, the evolutionary history of these worms is complex, with putative gene flow among ancestors. In this study, we investigated the early evolution of the family in relation to the diversification of thermal niches. We demonstrated that phylogenetic histories of alvinellid species greatly vary along chromosomes, possibly due to allele introgression between nascent ecotypes. We performed sequence reconstructions of ancestral cytosolic MDH and Cu/Zn SOD under the two most frequent phylogenies encountered along the genome. *In silico* simulations of folding stability for these two enzymes were highly correlated with their biophysical characterization (micro-calorimetry and differential scanning fluorimetry), and were used to predict the thermostability of thousands of alternatively reconstructed proteins. We further generalized the reconstruction at the proteome scale, taking the amino-acid usage bias as a proxy for folding stability in a new phylogenetic model accounting for amino-acid variations over time. Both approaches agree that the last common ancestors of Alvinellidae gained highly stable proteins, comparable to proteins of present-day thermotolerant species. Our computational protocol allowed us to validate this scenario under multiple phylogenetic hypotheses and different sequence reconstruction models. Considered together, Alvinellidae shed light on how metazoan species diversify in order to colonize different thermal niches, combining adaptive mutation and selection on genetic variants.

**Significance:** Deep-sea hydrothermal vents are among the most extreme environments on Earth. They, however, represent oases of life for a restricted number of highly specialized species. As such, Alvinellid worms constitute an exceptional family, including highly thermotolerant animals such as the Pompeii worm. Despite their vent endemicity, Alvinellidae thrive under contrasting ecological conditions, particularly regarding temperature. Understanding how this family diversified to colonize broad thermal regimes is of major importance for dissecting the mechanisms by which species adapt in highly unstable environments. Combining experimental protein resurrection, proteome-wide predictions, and new phylogenetic models, we assess the reliability of ancestral sequence reconstructions that represent molecular thermometers to evaluate past environmental conditions.

## 1 Introduction

In 1977, the submersible Alvin discovered the first deep-sea hydrothermal vents on the Galapagos Rift, at a depth of 2,500 meters [1]. These are among the harshest chemical environments known on Earth, yet highly specialized, chemosynthesis-based communities can thrive using the reduced compounds present in the hydrothermal fluid [2]. Animals inhabit the fluid mixing zone where they withstand extreme gradients of temperature and significant chemical variations in pH, oxygen, CO_2_, hydrogen sulfide concentrations [3]. Alvinellidae form one of the most ecologically diversified families of metazoan endemic to the vents. In contrast to other vent species, these polychaete annelids live closest to the undiluted fluid, on the walls of end-member chimneys, and experience regular temperature spikes, chronic hypoxia and high oxidative stress [4, 5]. Fourteen species are currently described, which have colonised along ridge systems of the entire Pacific and Indian Oceans [6, 7]. Although *in situ* measuring the unstable fluid chemistry remains challenging [8–10], Alvinellidae display striking adaptive features (e.g. hypertrophied gills, diversified hemoglobins, bacterial associations) linked to highly temporally variable thermo-chemical regimes they experience [9]. Amongst them, *Alvinella pompejana* is one of the most thermotolerant metazoans described to date, with a thermal optimum around 42 °C in pressurized aquarium [11]. Other species, such as the ecologically similar tube-builder *Paralvinella sulfincola*, also appear to have a thermal preference above 40 °C [12]. This is consistent with a previous work on the thermotolerance of *A. pompejana* and *A. caudata* mitochondria, which cease to function at 48-49 °C, compared to 31-33 °C for other deep-sea or shallow-water species living in cooler conditions [13]. Conversely, other Alvinellidae species such as *Paralvinella palmiformis* and *Paralvinella grasslei* are both mesophilic and avoid temperatures above 30 °C [12, 14]. Molecular studies also showed that the thermal activity range and folding stability of several enzymes are higher in ‘hot’-adapted species (*A. pompejana*, *A. caudata*, *P. sulfincola* and *P. hessleri*) compared to ‘cold’-adapted species (*P. palmiformis*, *P. grasslei*, *P. p. pandorae* and *P. p. irlandei*) [15, 16].

Therefore, there is a body of evidence suggesting that alvinellid worms tolerate a wide range of environmental temperatures and withstand extreme ecological conditions rarely encountered by any other metazoan species. How did these animals diversify in response to such contrasted environments is still an unresolved question. Based on *in silico* reconstruction of protein fragments from six species, it was suggested that the family descends from a thermophilic ancestor with some secondary selective relaxation in current colder lineages [17]. Yet, recent studies hint toward a complex history of speciation between the forming lineages, involving incomplete lineage sorting and potential gene exchanges, due to a rapid colonization of the vent habitat at the end of the Cretaceous period [5, 18].

In this study, we propose to validate the thermophilic state of the alvinellid worms’ ancestor and to better understand how the thermotolerance evolved at a molecular level according to the thermal habitat of these animals using phylogenetic and ancestral sequence reconstruction (ASR) [19], protein resurrection [20,21] and protein stability modeling [22–24]. Firstly, we investigated how the evolutionary history of genes might differ across the genome and reflect the adaptive radiation of these species in response to the rapid colonisation of thermally diverse hydrothermal habitats. Considering the most frequent alternative phylogenetic hypotheses that prevail along the Pompeii worm’s genome, we secondly experimentally measured the stability of enzymes on a large set of modern species as well as their probable reconstructed ancestors. We focused on the cytosolic malate dehydrogenase (cMDH) and the Cu/Zn superoxide dismutase (SOD). The cMDH is a homodimeric enzyme which activity is a reliable indicator of the thermal preference of marine invertebrates, including *A. caudata* and *A. pompejana* [25, 26]. The Cu/Zn SOD is primarily involved in the redox regulation, and plays a major role in the oxic/anoxic stress response of *P. grasslei* [27] and *A. pompejana* [28] when the temperature of the worm’s habitat varies. The experimental characterization was enriched with *in silico* predictions of the folding stability of thousands of alternatively reconstructed proteins at these two enzymes. Finally, we developed a new computational strategy relying on the amino-acid usage bias of proteins between ‘hot’ and ‘cold’ species to scale the method at the entire proteome level.

By combining biochemical analyses on resurrected proteins with a new in silico ancestral sequence reconstruction (ASR) approach at the proteome scale, we therefore advance our understanding of the molecular mechanisms by which ectothermic animals can evolve to survive in harsh and highly variable extreme thermal environments, and of how ecological specialization can promote species diversification in metazoans.

## 2 Materials and Methods

### Identification of orthologous genes

We used transcriptome assemblies previously obtained by [18] from RNAseq data of Alvinellidae (12 species) and outgroup ampharetid and terebellid species (*Hypania invalida*, *Amphicteis gunneri*, *Neoamphitrite edwardsii*, *Streblosoma kaia*, *Melinna palmata*, *Amphysamitha carldarei*, *Anobothrus* sp.), and the Pectinariid *Pectinaria gouldii* retaining all possible transcripts longer than 200 nucleotides. We also performed a partial genome assembly for *P. pandorae irlandei* using a illumina sequencing and SPades [29], gene prediction using Augustus [30] and *de novo* assembly of RNAseqs reads from Fontanillas and colleagues [17] using Trinity v. 2.9.1 [31] to increase the quality of coding sequences for this species. Details and scripts are given in supplementary information.

Orthologous transcripts from all polychaete RNAseq data were determined by reciprocal best-hit blastx against the complete gene catalog obtained from the *A. pompejana* genome [32]. We also added orthologous groups identified by [18], either complete sequences or sequences longer than 120 amino acids, containing one *P. p. irlandei* sequence. Finally, orthologous genes were aligned using MAFFT L-INS-i [33], and fragments smaller than 20 amino acids between two gaps were considered unreliable and removed from the alignments. If orthologous groups obtained from this study or from the initial analysis of [18] pointed to the same gene, we only kept the alignment containing the longest transcripts.

### Gene coalescence mapping along the 16 chromosomes of the genome of *A. pompejana*

Brun and colleagues [18] identified 15 potential gene topologies at the most ancestral node of the alvinellid worms. Using IQ-TREE (LG+G+F model) [34], we tested these topologies with minimal discriminative groups of species to reduce potential uncertainty at other nodes in the trees. The discriminative groups contain only *A. pompejana*, *P. unidentata*, *P. pandorae irlandei*, one or two other *Paralvinella* species, one or two Ampharetidae, one or two Terebellidae (see supplementary figure 1). Each gene was associated with its most likely topology out of the 15 candidate phylogenies, and mapped to their corresponding locus onto the *A. pompejana* genome.

The unequal distribution of the most frequent T6 and T9 distributions were tested by two different tests at a 2.5% p-value threshold to take into account a multi-test correction at 5%. The first test’s null hypothesis is that the observed distribution of genes belonging to *T* over genomic windows falls within the expected fluctuations of *T* frequencies, and aims at determining whether *T* is over-represented in one particular window of the genome. The second test’ hypothesis is that *T* is equally interspaced over the genome. By aggregating the weaker signals across all genomic windows, this test sacrifices window-level resolution to determine whether *T* is preferentially concentrated in specific genomic regions rather than uniformly distributed across the genome. The two tests rely on expectations obtained from 5000 simulated genomes where the gene topology and localization are randomly shuffled, and the over/under-representation of a topology is tested *via* binomial tests (*B_T_* (*n, p*)), where *n* is the number of genes found in a genomic window, and *p* is the empirical frequency of a given topology *T* over all genes. Details are provided in supplementary figures 4A and 4B. We consider 2.5 Mb windows sliding every 250 kb along the chromosomes (1231 windows with 48.5 genes on average), which is smaller than the length of the detected exchanged window.

### Overexpression of modern and ancestral cytosolic malate dehydrogenases and Cu/Zn superoxide dismutases

We chose to focus on two enzymes: the cytosolic malate dehydrogenase (cMDH) and the Cu/Zn superoxyde dismutase (SOD) in order to get two independent information on the putative thermal habitats of the ancestors. The complete cMDH and Cu/Zn SOD transcripts were retrieved from the transcriptomes of the 12 modern alvinellid species and the outgroup species. The ancestral sequence reconstructions (ASR) were performed under the two most likely phylogenetic hypotheses T6 and T9. Ancestral sequences corresponding to maximum likelihood estimates were computed under the LG+G4+F model with branch optimization, equivalent to a classical ASR procedure, using the models M6 and M9 (see paragraph *Reconstruction of Alvinellid ancestral proteins at the proteome scale*) to test the effect of incorporating amino acid bias variation in the reconstructions.

We chose to express the recombinant proteins of modern thermophilic species *A. pompejana*, *P. sulfincola*, *P. fijiensis*, *P. mira*, and the cold-adapted species *P. grasslei*, *P. palmiformis*, *P. p. irlandei* and *P. unidentata*, as well as ML estimates of ancestral proteins corresponding to all internal nodes of the Alvinellidae tree, excluding recent ancestors of pairs of species sharing similar thermal habitats. To this end, competent *E. coli* were transformed with pET100/D-TOPO expression vectors (ThermoFisher) containing the retro-translated protein coding sequence fused with a (His)6 tag. Proteins were then purified on *Ni*^2+^ chelation chromatography followed by size-exclusion chromatography with a HiLoad 16/60 Superdex 75 (Cytiva) column in 20 mM Tris-HCl, 200 mM NaCl pH 7.4 buffer. Details of the expression procedure are given in supplementary material.

Thermal unfolding of the cMDH was measured by nano Differential Scanning Fluorimetry (nanoDSF) using the Prometheus NT.48 instrument (NanoTemper). Samples were heated from 20 °C to 95 °C with a temperature increase of 1 °C/min. The fraction of unfolded proteins *f_u_*(*T*) was derived from the 350/330 nm fluorescence ratio *r*(*T*), assuming a linear relationship between the signal and the unfolding fraction. Data were fitted against the temperature *T* in Kelvin using the apparent Gibbs-Helmholtz model [35], see supplementary material:

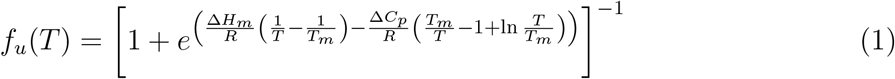

where *R* is the molar gas constant, *T_m_* is the melting temperature at which half of the proteins are denatured, Δ*H_m_*the enthalpy variation at *T_m_*, and Δ*C_p_* the heat capacity change of the protein during unfolding. These parameters were all estimated from the *in vitro* measurements of the protein denaturation.

Thermodynamic parameters for the Cu/Zn SOD were measured by Differential Scanning Mmicro-Calorimetry using the Microcal PEAQ-DSC (Malvern), as these proteins do not contain tryptophanyl residue to monitor their fluorescence with nanoDSF. The excess heat capacity 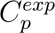 was measured between 20-110°C, with a temperature increase of 1 °C/min. 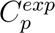 is fitted against the temperature according to [36] (see supplementary material)

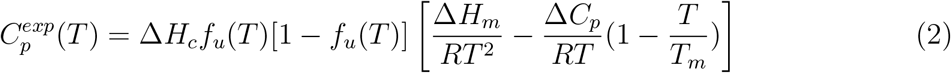

with *f_u_*(*T*) being expressed as in equation 1. We considered three potential successive denaturation events for the SOD [37]. The measured signal is consequently assumed to be the sum of three distinct 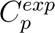, with their own thermodynamic parameters to estimate. The third denaturation event is assumed to correspond to the unfolding of the most stable holo-enzyme, while the other events correspond to structures that have lost their Cu or Zn ions.

### Estimating confidence envelopes of the ancestral reconstructions of the two enzymes

In order to assess the stability of ancestral proteins at a given node when taking into account the uncertainty of ancestral residues in the reconstruction, 1000 alternative copies of the reconstructed proteins were sampled from the posterior sequence distribution of the residues. Marginal probabilities of residues above 0.95 were rounded to 1 to avoid the introduction of numerous mutations [38]. The stabilities of the alternative proteins were predicted using FoldX [23] and the wild type reference protein structure of the two enzyme families obtained with Colabfold2 [39]. The predicted stabilities of FoldX were then converted into potential experimental stabilities using the linear regression:

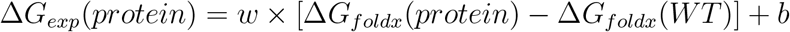

The linear coefficients of the equation were estimated both for the cMDH and Cu/Zn SOD from the Δ*G_exp_*of experimentally characterized proteins and their Δ*G_foldx_* stability predictions. Scripts for this procedure are available on the GitHub repository.

### Reconstruction of Alvinellid ancestral proteins at the proteome scale

In ancestral sequence reconstruction, classical models consider the past amino acid frequencies as constant over time and equal to the mean frequencies in modern species [40–42]. However, amino acid usage is not expected to remain constant, and could especially be influenced by environmental temperature [22, 24, 43–45]. We therefore developed an evolutionary model where the equilibrium frequencies of amino acids are allowed to vary along the tree branches.

To achieve this, we decompose the average amino acid frequencies in proteins of modern species with respect to their principal components (PCA). In this decomposition, amino acid frequencies in species *s* can be written as 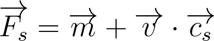

where 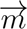 is the mean frequencies of amino acid across all species, *v* is the PCA eigenvectors and 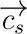 is the coordinates of the species *s* in the PCA space.

In our reconstructions, we introduced the coordinates *c* for the ancestral species as well, which estimates are found by maximum likelihood optimization. To reduce the number of coordinates (and therefore of parameters) of the model, we reduced the PCA space to only its first largest components to approximate amino acid frequencies. The number of PCA to retain is determined by the AIC criterion, which balances the likelihood increase of the model with a penalty on the number of parameters.

Along branches, sequences evolve under a hidden Markov Model, with a probability kernel obtained as *M* = *F_s_ × Q*, with *Q* being chosen as the LG matrix. In our implementation, a model with no eigenvector is exactly equivalent to a LG+G4+F model. Details on the method and algorithms are given in supplementary material and on the Github repository.

The parameters of fifteen evolutionary models, M1 to M15, were learned independently on concatenation of genes strongly associated with each of the fifteen topologies (*Pr >* 0.5). Site positions from which more than two species were missing were discarded beforehand from the sequence alignments. To estimate the robustness of our results under different topological hypotheses, we used the optimized models M1 to M15 to perform marginal ancestral sequence reconstruction on an independent set of translated genes that did not match exactly any of the 15 topologies. For these alignments no missing species was allowed at any residue position of the alignment. The ancestral amino acid bias was reconstructed from these genes and subsequently used to delimit envelopes of uncertainty around the most important ancestral nodes due to the different phylogenetic histories between genes.

### Evolution of protein thermostability indices along the Alvinellid lineage

The buried positions were extracted based on secondary structure predictions for *A. pompejana* proteins with Netsurp 3.0 [46]. In full sequence alignments, we consider that the protein folds are well conserved. As a consequence, residues that align to the buried positions of *A. pompejana* proteins are buried as well in the other species. The amino-acid compositions of buried sites were finally displayed graphically alongside axes expected to reflect protein stability according to Tsuboyama and colleagues (see figures 3a and 3b of the aforementioned article) [47].

## 3 Results

### A Genome-wide discrepancy of gene histories: the consequence of a rapid radiation of Alvinellidae due to habitat diversification?

Brun and colleagues previously identified incongruent gene histories in alvinellid species, based on a set of 699 orthologous transcripts, and attributed these incongruences to a high level of incomplete lineage sorting (ILS), as well as some potential selective cross-species gene exchanges (CSGE) during the rapid radiation of this worm family [18]. We firstly described more precisely how genes evolved during the birth of alvinellid worms using a larger set of 6639 gene transcripts, mapped on a recently published genome assembly of the *A. pompejana* at a chromosome level [32]. The two previously identified topologies T6 and T9, in which *P. unidentata* is sister species with either *P. p. irlandei* or with the *Alvinella* species, were confirmed to be the most represented in the genome (respectively 10.2 and 8.7% of the genes, supplementary figure 2 and 5) but formed a mosaic with other topologies all along the 16 chromosomes.

Localising gene topologies onto the genome did not support our initial hypothesis that some genomic regions could display specific histories due to differential selective pressures (see figure 1). We tested whether 2.5 Mb genomic windows sliding every 250 kb contained an excess of the T6 or T9 using the empirical genome-wide frequencies of gene tree topologies. The null hypothesis was that if the gene topologies are randomly distributed along chromosomes, no genomic window can contain an excess of one of these topologies. Only one window, spanning 300 kb on chromosome 4, exhibited a significant over-representation of T9 (p-value 4.2% after multi-test correction; see Supplementary Figure 4.A), which is consistent with a trans-specific selective sweep. However, this alone was not enough to explain the previously observed global imbalance of gene histories in the transcript pool. Considering the genome’s overall organisation, the T9 and T6 topologies, although preponderant, are interspaced with other topologies across the genome in a similar way to that observed in simulated genomes with randomly shuffled gene localisation (although one minor topology T10 is detected as marginally significant, see supplementary figure 4.B).

**Figure 1:**
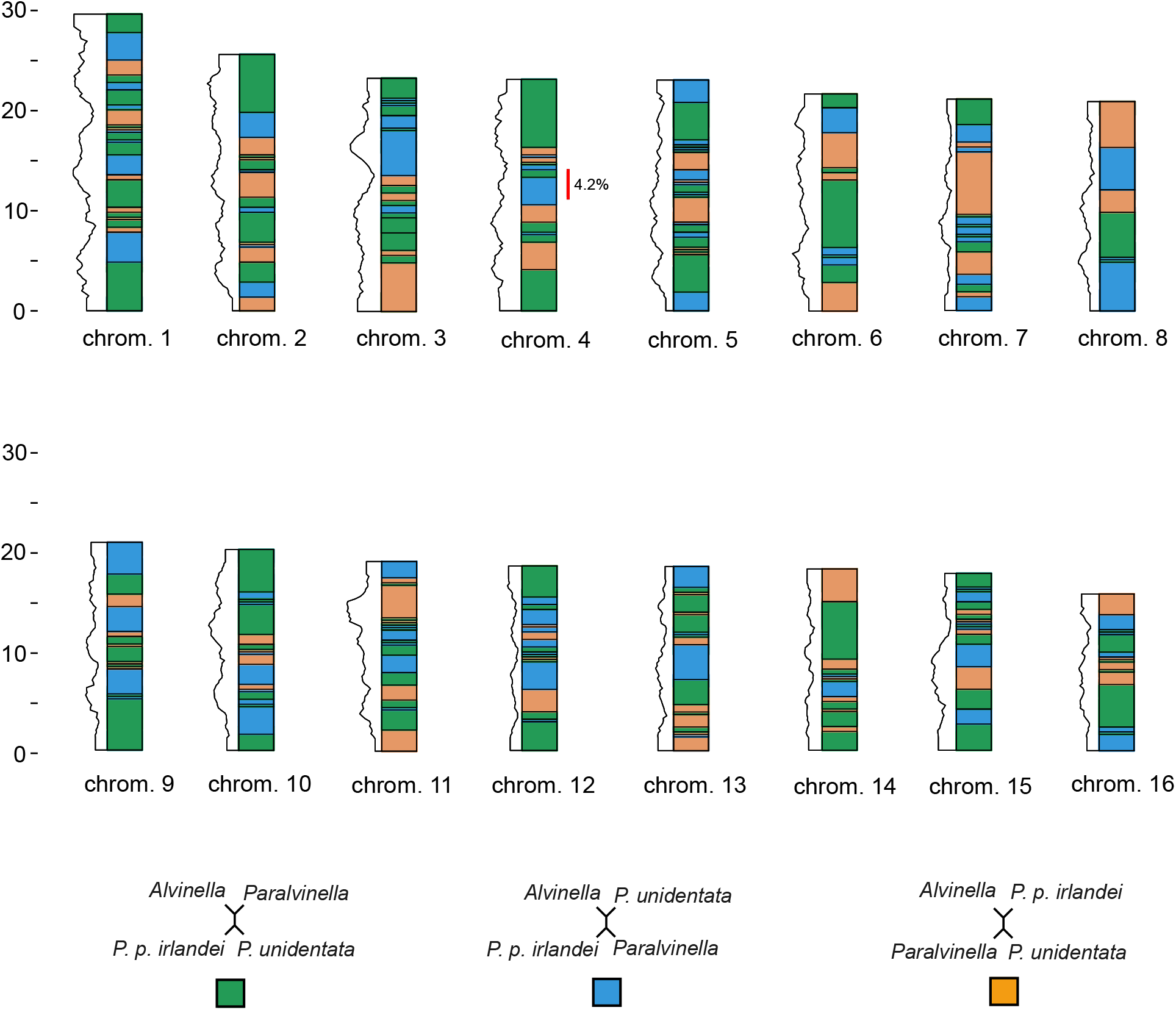
Gene topologies mapping along chromosomes of *A. pompejana*. The rearrangement of species is likely to be the result of specific gene histories associated with incomplete lineage sorting, cross-species gene flow. The chromosome length is given in Mb. Using a 2.5 MB window, sliding every 0.25 MB, we observe a mosaic of regions along the genome where one gene topology is more represented than the others. Only one region of chromosome 4 displays a higher-than-expected proportion of gene histories bringing the *P. unidentata* and the *A. pompejana* lineages together with a p-value of 4.2%.

The alternation of gene topologies along the AP genome (see supplementary figure 5) without any clear architectural patterns suggests that either ILS or neutral CSGE occurred on a large scale during the family’s radiation. This pattern, observed at all degrees of confidence filtering of the gene topologies, may have been driven by an extremely rapid species diversification within a confined spatial area where inter-population exchange was still possible, and by the long-term maintenance of cross-species allelic lineages due to adaptive combinatorial mechanism of genetic variants [48].

The alternation of gene topologies along the AP genome (see supplementary fig. 4), without any discernible architectural pattern, suggests pervasive ILS and/or CSGE during the family’s radiation. Importantly, this pattern remained unchanged after progressively restricting the analysis to genes with increasingly well-supported topologies, indicating that gene tree estimation error is unlikely to explain the apparent genomic randomness. This is consistent with an exceptionally rapid species diversification within a geographically restricted area, where gene flow among emerging populations remained possible, together with the long-term maintenance of allelic lineages across species through adaptive combinations of genetic variants.

Following this result, the fundamental imbalance between gene topologies was accounted for in subsequent steps of ancestral protein reconstruction (ASR) in order to strengthen our conclusion about the thermophilic nature of alvinellid ancestors.

### The last common alvinellid ancestor evolved toward thermotolerance, but not thermophily

In order to assess the ancestral thermal habitat of alvinellids and how ancestors evolved toward thermophily in some modern lineages, we chose to perform ancestral sequence reconstruction of the cytosolic malate dehydrogenase (cMDH) and the Cu/Zn superoxide dismutase (Cu/Zn SOD) under both the T6 and T9 topologies, which were confirmed to be the most frequent gene histories genome-wide. Six ancestral proteins per enzyme and topology were reconstructed by maximum likelihood at key nodes of the species phylogeny, where potential ‘hot’ versus ‘cold’ environmental transitions could have occurred. The folding stability of these resurrected enzymes were later experimentally assessed after being expressed in *E. coli* together with the modern ones.

We measured the unfolding thermodynamic parameters of the expressed recombinant enzymes *in vitro* on a ramp of temperatures (see supplementary figure 5 and table 5, 6). As shown in figure 2, the melting temperature (*T_m_*, at which half of the protein is denatured) of modern species was narrow but sufficient to separate ‘hot’-adapted from ‘cold’-adapted species. Under the T6 topology hypothesis, the melting temperature of the last common ancestor of Alvinellidae (Anc1) was by one or two degrees higher than under the T9 topology. Moreover, for the cMDH, ancestral *T_m_* were between 60 and 62 °C, which lies in the low range of modern thermophilic species (*P. mira*, 61 °C). Because the average of *T_m_* differ between cMDH (56 to 66 °C) and Cu/Zn SOD (72 to 82 °C) by more than 15 °C, the temperature difference between ‘hot’-adapted and ‘cold’-adapted species was even more pronounced for the Cu/Zn SOD, with ancestral protein *T_m_* ranging between 78 and 83 °C. They were more closely-related to the ‘hot’-adapted species (*P. fijiensis* and *A. pompejana*: 80 °C) when compared to the ‘cold’-adapted species (*P. unidentata*: 76 °C). These results clearly indicated that the oldest alvinellid ancestors were mesophilic to mildly thermophilic.

**Figure 2:**
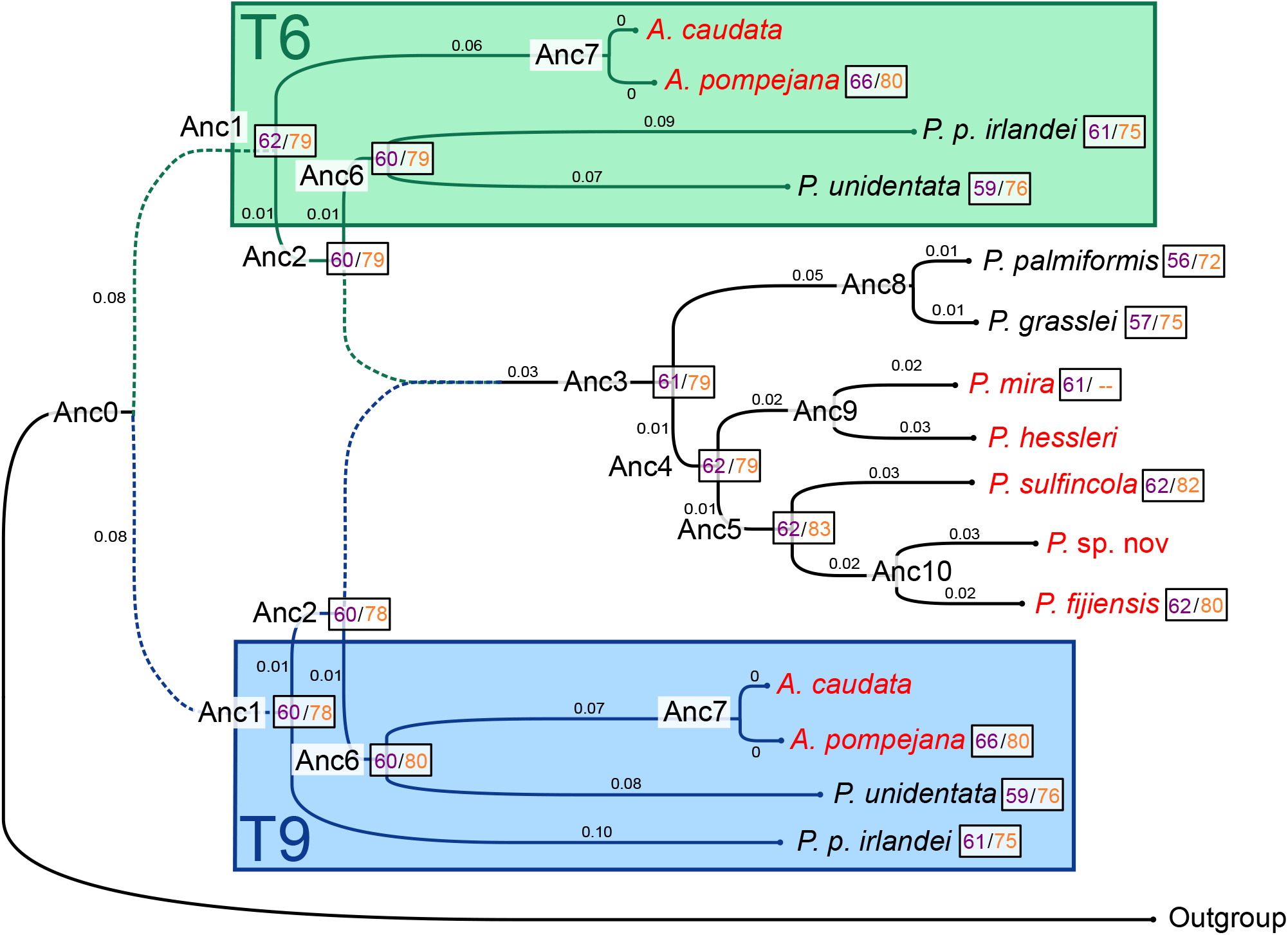
Melting temperatures for modern and putative ML ancestral reconstructions of cMDH (first value in purple) and Cu/Zn SOD (second value in red) under the two main phylogenetic hypotheses (T6 and T9). Branch lengths correspond to expected number of amino acid replacements per residue according to [18]. Modern hot-adapted species are written in red.

As reconstructed ancestral proteins can show artificially high stability, especially when only considering the maximum-likelihood sequence [49–51], we predicted the stability of other alternative proteins, sampled from the posterior distribution of the reconstructed sequences [49]. To this end, we predicted the stability of 1000 alternative sequences for each ancestral node of the Alvinellidae phylogeny using either T9 or T6. As shown in figure 3A and B, the FoldX [23] predicted protein stabilities for both the cMDH and Cu/Zn SOD were strongly correlated with the experimental measurements (*R*^2^ = 0.93 and 0.84, respectively), after the removal of duplicated maximum likelihood sequences and two predictions of *P. fijiensis* cMDH and Anc3-T6 Cu/Zn SOD that were inconsistent with the experimental measures. The regressions were later used to simulate the experimental ΔΔ*G* of ancestral proteins of the family, while accounting for sequence reconstruction uncertainty.

**Figure 3:**
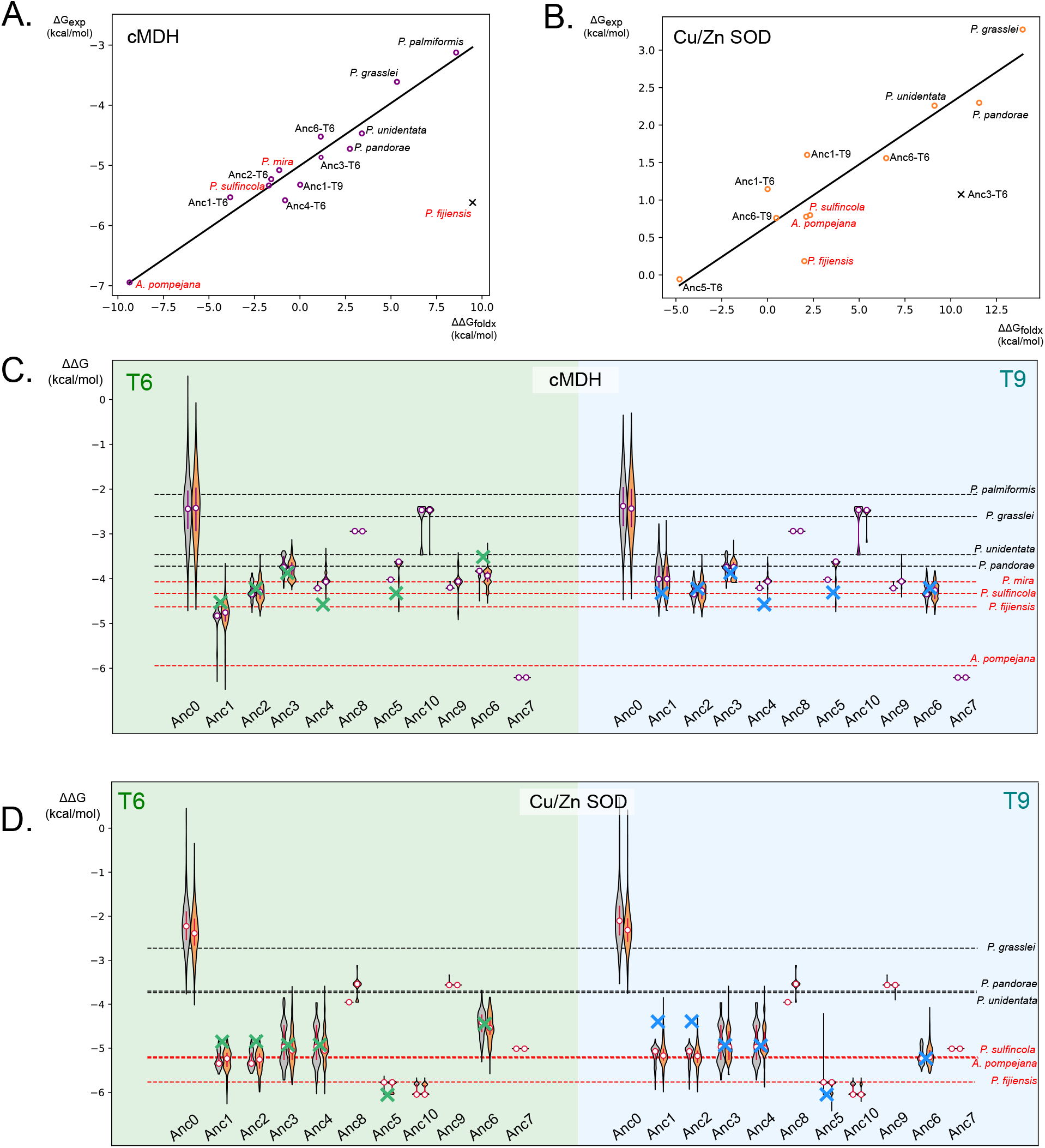
Ancestral protein reconstructions of the cMDH and Cu/Zn SOD. On all panels, modern hot-adapted species are written in red. A/B. Correlations between the experimental and FoldX predicted stabilities of cMDH and Cu/Zn SOD, respectively. Dots marked with a ‘x’ are considered as outliers, and excluded from the correlation. C/D. Ancestral protein reconstructions of cMDH and Cu/Zn SOD. Experimentally measured protein stabilities of modern species and of the maximum likelihood ancestral sequences Anc1 to 6 are in dashed lines and marked by a cross, respectively. The envelopes of confidence correspond to *in silico* predictions obtained by FoldX on 1000 alternative proteins at each node of the phylogeny. The alternative reconstructions are either obtained with the LG+G4+F model (grey), or with an evolutionary model accounting for amino-^1^a^5^cid composition variation over time (brown), with parameters estimated onsets of alvinellid orthologous genes congruent with topologies T6 and T9. Medians and interquartile ranges are displayed.

The experimental measures of the protein stabilities of extant species allowed us to match protein stabilities with expected environmental temperatures. Modern ‘hot’-adapted and ‘cold’-adapted species in the two protein families are indeed clearly separated (Figure 3C and D). We observed that the last common ancestor (LCA) of Ampharetidae and Alvinellidae (Anc0) had protein stabilities similar to those of *P. grasslei* and *P. palmiformis*, followed by a clear increase of the stability toward the alvinellid LCA (Anc1) for the two phylogenetic hypotheses. This suggests that the initial split leading to the emergence of the Alvinellidae family occurred under already mesophilic conditions, followed by a thermal regime increase in the alvinellid lineage. Accounting for amino-acid composition fluctuations in the evolutionary model (brown envelopes in panels C and D) has only a negligible impact on the stability distribution of proteins and leads to the same conclusions.

Surprisingly, the Anc10 cMDH and Anc9 SOD have predicted protein stabilities closer to ‘cold’-adapted modern species. The direct descendants of these nodes (cMDH *P.* sp. nov, *P. fijiensis*, or SOD *P. mira* and *P. hessleri*), despite being ‘hot’-adaptedspecies, were also predicted with a low protein stability by FoldX (see supplementary table 8). These two ancestral sequences likely inherit common mutations that are wrongly inferred as less stable by FoldX. This unexpected and contradictory result implies that *in silico* stability predictions of enzymes in Anc0, Anc7, Anc8, and Anc10) have to be taken with caution, as no variant was tested experimentally.

Nevertheless, the experimental characterization of protein thermostability of these two enzymes, in accordance with computational predictions, validates our view that the ancestor of Alvinellidae rapidly acquired new mutations to live at higher temperatures after the split from the common ancestor of Alvinellidae and Ampharetidae, which was pre-adated to moderate temperatures. This initial proteome adaptation was then followed by some relaxation of the protein stability in lineages leading to the ‘cold’-adapted modern *P. unidentata*, *P. pandorae*, *P. grasslei* and *P. palmiformis* while protein thermostability was reinforced in ‘hot’-adapted modern species.

### The proteomes of Alvinellidae show a strong adaptive signal toward protein thermostablity

On a broader scale, we examined how temperature changes in the Alvinellidae habitat could influence the molecular evolution of proteins at the proteome level, and how this evolution could, in turn, have promoted the ecological species diversification of these worms. We therefore investigated the species adaptation to temperature by analysing hundred orthologous proteins of all the modern and ancestral species of alvinellid, ampharetid and terebellid groups. As can be seen in figure 4A., the amino acid usage in Alvinellidae is specifically biased towards E, L, Y, K and A, and depleted in D, G, M, S and Q, compared to closely related coastal but also deep-sea terebelliform species. This biased amino acid usage has already been observed between six ‘hot’and ‘cold’-adapted alvinellid species [17], and has been suggested to be an adaptive trait for living in higher environmental temperatures.

**Figure 4:**
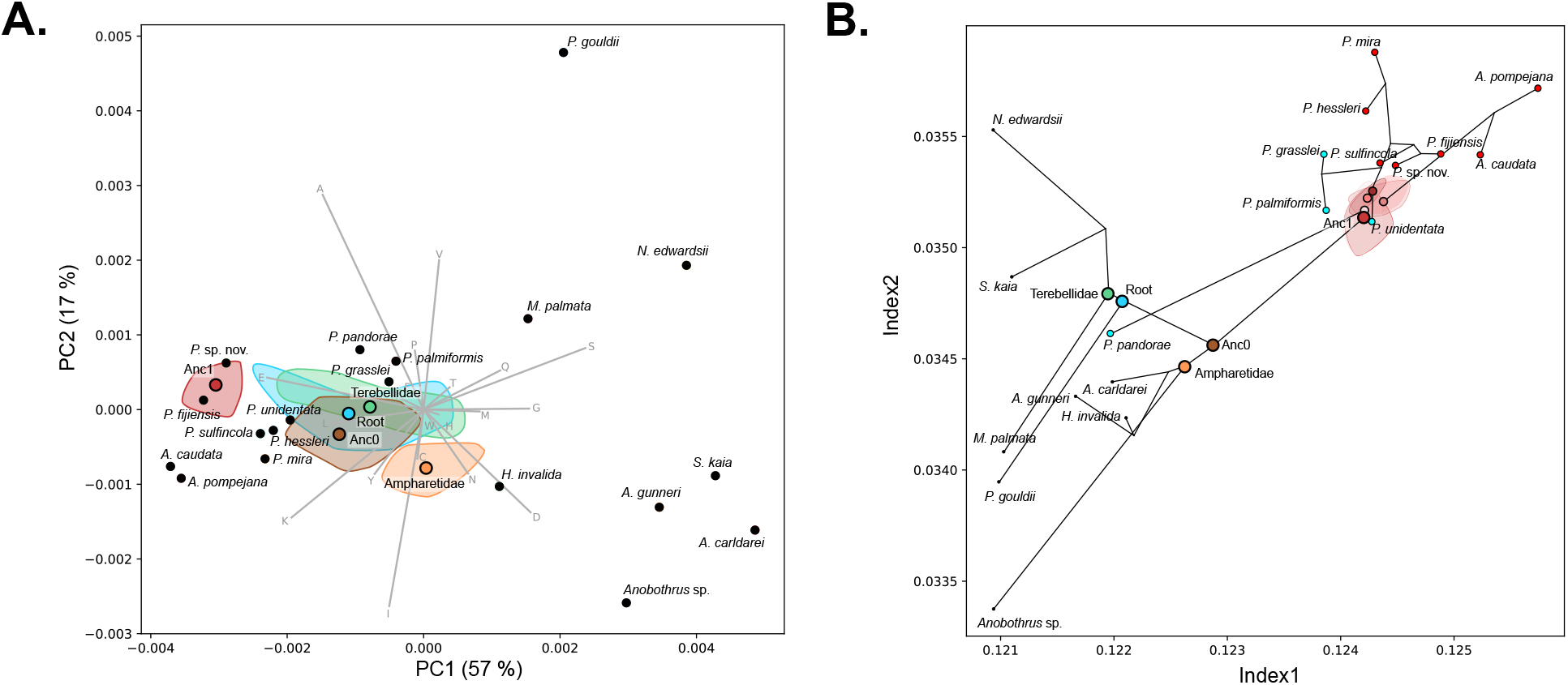
Amino acid usage bias estimated from 263 proteins in the alvinellid family. For the ancestral reconstructions, the uncertainty envelopes due to the different phylogenetic hypotheses are displayed, and the mean coordinates are weighted by the expected number of genes falling under each of these hypotheses. A. Principal components analysis on the amino acid usage in alvinellid species, their ancestors and other deep-sea or coastal terebelliform polychaetes. B. Projection of the buried amino-acid compositions of ancestral and modern alvinellid species with respect to the two indexes proposed by Tsuboyama and colleagues [47]. These indexes are expected to separate the most thermostable proteins (top right) from the least stable ones (bottom left).

The biased amino acid usage prompted the development of an evolutionary framework that accommodates variation in amino acid frequencies in proteins through time (see material & methods and supplementary material and supplementary figure 7, 8). We derived 15 phylogenetic evolutionary models with maximum likelihood parameters, each corresponding to one of the 15 plausible gene histories. The models were optimized using sets of concatenated genes that were each strongly associated with a peculiar gene history (posterior probability *>* 0.5). Ancestral reconstructions, accounting for amino acid usage bias, were then tested on another set of 263 complete transcripts for which we could retrieve proteins from all the species. It appears that the alvinellid ancestors had amino acid compositions similar to those of *P. fijiensis* and *P.* sp. nov. on a 2D PCA projection (74% of variance extracted by the first two axes, see figure 4A.). This is very different from the last common ancestor between Alvinellidae and Ampharetidae, whose composition is closer to that of other terebellid ancestors. This suggested that a molecular signature specific to Alvinellidae worms was gained during the initial step of the vent colonization.

Tsuboyama and colleagues [47] were able to correlate the fold stability of small protein domains with preferred amino-acid replacements. Over the whole experiment, 46% of the observed stability was explained by two linear combinations of amino-acid frequencies, in which the gain of large, non polar or aromatic residues is likely to stabilize the core of proteins. We tried to consider these linear combinations to construct two indices of folding stability, named Index1 and Index2 hereafter. These indices were used to score the expected relative stability of domains from the 263 proteins in our modern and ancestral species. The domains were predicted considering the fold of *A. pompejana* proteins as a reference for all other aligned proteins of species. The projection of the domain’s stabilities according to these two indices was not able to differentiate ‘hot’-adapted and ‘cold’-adapted species when taking all the amino acid residues into account (see supplementary figure 3). However, predicted buried residues were highly correlated with the expected protein stabilities of modern species. ‘Hot’-adapted species have indeed the largest indices on both Index1 and Index2 (see figure 4B).

The colonization of the vent habitat by alvinellid ancestors is also visible along the evolution of the two thermostability indices depicted by Index1 and Index2. Different phylogenetic hypotheses introduced variations in the stability of the reconstructed proteins, as shown by the different envelopes of uncertainty at the birth of modern lineages. However all estimations overlap in the same region of the graph and we expect that alvinellid ancestral populations experienced similar temperature adaptation during the colonization of the Pacific and Indian ridges. Figure 4B also reinforces our view that modern alvinellid species have undergone parallel evolution by independently increasing or reducing their thermotolerance when colonizing different thermal niches (‘hot’ chimneys for *Alvinella* or *the P. mira* and *P. hessleri* lineages, or ‘cold’ diffuse venting for *P. palmiformis* or, strikingly, *P. p. pandorae*.

The habitat shift of the first Alvinellidae ancestors is clearly associated with the emergence of a strong adaptive signature at the proteome scale. These lineages were likely sharing the same thermal regimes despite phenotypic variation, rapidly specializing to the hottest and coldest parts of hydrothermal vents until they became geographically separated to occupy non overlapping areas nowadays.

## 4 Discussion

### A long history of molecular adaptation and diversification in the hydrothermal environment

For proteins evolve toward marginal stability at the intracellular temperature [52, 53], ectothermic organisms are expected to show a positive correlation between environmental temperature and protein stability [54]. Nevertheless, most Alvinellidae exhibit eurythermal behavior *in situ* and can tolerate prolonged temperature fluctuations, hidden within parchment-like tubes or mucus cocoons [12, 14]. Our measurements across two enzyme families were however in line with the assumption that modern alvinellid worms can have different thermal *preferenda* [12, 15] with a clear signal of molecular adaptation to distinct thermal regimes. In particular, the folding free energies (Δ*G*) distinguish proteins from ‘hot’and ‘cold’-adapted lineages.

The resurrection and experimental characterization of ancestral cMDH and Cu/Zn SOD demonstrated a clear increase in the thermal *preferendum* of the alvinellid LCA with the introduction of mutations stabilizing the protein folds. These biophysical measures were combined with computational predictions of thousands of reconstructed variants that produced robust envelopes of confidence around the ML sequences. Most of these envelopes were narrow, with a range of stability values similar to those found in modern ‘hot’-adapted species. Finally, our new amino-acid based computational approach produced exactly the same conclusions regarding protein stability, while scaling the method to a large number of proteins in the context of the species’ complex evolutionary history. Thermostability indices [47] computed for 263 ancestral proteins agreed with the experimental enzyme data, indicating a progressive increase in proteome thermostability toward the mildly thermophilic Alvinellidae LCA. This ancestor was likely similar to modern species such as *P. sulfincola* and *P. fijiensis* but not as thermophilic as specific ‘hot’-adapted alvinellid lineages such as the *Alvinella* lineage. The subsequent diversification of the family into distinct vent habitats produced protein repertoires with only ̂10% average sequence divergence that nevertheless remain adapted to contrasting thermal regimes. This adaptation appears to rely primarily on substitutions in protein cores rather than at solvent-exposed residues, which experience different evolutionary constraints [55, 56].

According to the last recent estimates, the radiation of Alvinellidae occurred between 80 and 110 Mya [18] in the Eastern Pacific before the emergence of the East Pacific Rise due to the subduction of the Farallon plate under the American one [57]. Interestingly, the LCA between Alvinellidae and their sister-group Ampharetidae displays proteins that are as stable as those of the coldest current Alvinellidae (*P. grasseil* and *P. palmiformis*, and more stable than those of some other hydrothermal Terebellidae and Ampharetidae (*S. kaia* and *A. carldarei*). This may indicate that the ancestor to Ampharetidae and Alvinellidae either colonized deep-sea hydrothermal vents before the diversification of Alvinellidae and Ampharetidae, or, alternatively, was inhabiting globally warmer waters. Deep Pacific temperatures peaked during the Late Cretaceous greenhouse (100–75 Mya), reaching about 10–15 °C [58, 59], and then cooled down until *≈*65 Mya before rising again during the Paleocene–Eocene Thermal Maximum (56–47 Mya) [60–62]. Because these temperature maxima occurred well after the divergence of the Alvinellidae and Ampharetidae ancestors (110–160 Mya) [18], ocean warming alone cannot explain the high stability of these ancestral proteins. Then, thermophily likely arose in alvinellid ancestors during the colonization of ‘hot’ vents and was later independently strengthened in several lineages (*Alvinella*, *Paralvinella Miralvinella*) or lost in others (e.g., *P. palmiformis*, *P. grasslei*, *P. pandorae*), reflecting separate episodes of independent adaptations to distinct thermal habitats within hydrothermal environments.

### Footprint of past ILS and CSGE during the alvinellid radiation with possible selective combination of ‘old’ polymorphisms shared between species

Gene topology weighting discrepancies along the AP genome suggest that ancestral Alvinellidae populations experienced possible ILS and CSGE due to a rapid radiation [63–65]. The pattern of topological alternations, sliced into very small portions of chromosomes and across chromosomes overall, likely represents a fossil footprint of an old but rapid radiation event (a flock of species [66]). The older the radiation event, the smaller these regions are [67]. This fits well with the molecular dating of this event. However, at least one local significant over-representation of a specific genealogical history on chromosome 4 of the AP genome (as well as another global imbalance regarding a minor phylogenetic scenario) suggests the signature of ancient selective effects between emerging Alvinellidae lineages, with selection favoring trans-species exchanges of pre-adapted alleles [68, 69]. The combination of balanced diversifying selection and ILS, arising from the adaptive radiation of worms across contrasting thermal habitats, may also contribute to the discordance among gene genealogies along the genome, resulting in the observed mosaic of gene topologies [48, 70].

Moreover, admixture between hybridizing species can also accelerate adaptation to environmental changes by keeping foreign alleles that perform better locally [71] and provide rapid selective advantage [72–74]. This process enables divergent ecotypes to reuse standing genetic variation through hybridization during ecological transitions [75, 76].

In modern alvinellid species, adaptive polymorphisms have been previously identified with allozymes exhibiting different thermostabilities in *A. pompejana*, *P. sulfincola* or *P. palmiformis* [15, 77, 78]. Balanced selection of locally well-adapted variants is likely maintained at broader scales by the high spatial heterogeneity and temporal variability of the vents [70, 79]. This is especially the case of the gene that encodes the phosphoglucomutase 1, which displays two variants in the Pompeii worm *A. pompejana*, one ‘old’ lineage of thermostable alleles associated with the colonization of newly formed hot vent chimneys and one associated with older, colder chimneys [77, 80]. These ecotypes demonstrate the adaptive potential for the ecological segregation of populations, *prior* to speciation if gene flow is disrupted: a situation already encountered between the northern and southern populations of the worm along the EPR [81].

Together, ongoing gene exchange between differentially temperature-adapted ecotypes within modern species and our deep phylogenomic analyses support the view of an ecological diversification of this worm family through similar processes, in geothermally active habitats, where seafloor dynamics can alternately isolate or reconnect populations [82].

### Confidence in the reconstruction of ancestral enzymes and *in silico* predictions

ASR has already been used to infer past environmental conditions of extinct species [83, 84]. However, some authors have suggested that a consensus effect could favor the fittest aminoacids and artificially increase the stability of reconstructed proteins, especially when considering only the maximum likelihood sequences [21, 49–51]. Here, we show that combining experimental characterization of protein stability with computational predictions improves the robustness of ASR. The FoldX [23] predicted stability of cMDH and Cu/Zn SOD is highly correlated with the experimental measures in both modern species and ML ancestors. This relationship can be used to predict the stability of ancestral proteins for which experimental characterization is not achieved, as well as that of alternative ancestral sequence variants at given nodes of the phylogeny. The resulting distribution of predicted stabilities provides confidence intervals that account for uncertainty in ancestral state reconstruction, including the contribution of less likely, potentially suboptimal residues.

Although physical modeling has proven to be in good agreement with experimental measures, some proteins failed to be predicted correctly. This can be attributed to the difficult task of interpreting the effect of antagonistic mutations that not only alter the immediate interactions of residues with their surroundings, but also the compactness of the protein [85]. In our case, discrepancies between measurements and predictions for several extant ‘hot’adapted species allowed us to extrapolate the uncertainty associated with the prediction of the stability of their recent ancestors, because these species share most of the substitutions underlying the prediction error with their immediate ancestors.

Predicting the stability of proteins is a long-standing goal in biology, with methods relying on the physical modeling of the protein’s force fields such as FoldX, and others using machine learning algorithms [23, 86–88]. Yet the choice of protein families, folds, and residues is also critical when linking environmental temperature to sequence stability, as observed stability is also the outcome of several extrinsic factors such as misfolding avoidance with chaperones or ligands and stabilization with cellular osmolytes [56, 89–92]. One main technical conclusion of our study is that experimental measurements of protein stability must be conducted on many, if not all of the modern species to aggregate statistical signals and safely infer the thermal habitats of ancestors.

We therefore complemented these simulations with a simple inference framework based on amino-acid usage bias, which is both highly scalable and interpretable. To this end, we developed a new evolutionary model that incorporates the historical evolution of amino-acid usage, thereby capturing the variation in lineage-specific compositional signatures. This approach primarily affects reconstructions of deep ancestral sequences separated from extant reference taxa by long branches, where uncertainty is greatest, and is expected to be less sensitive to artefactual convergence arising from independent shifts in compositional heterogeneity across lineages [93, 94]. The algorithm is fast and the optimized model proved to better fit the observed data, according to the corrected Akaike Information Criterion.

## 5 Conclusion

The role of habitat diversification, and how environmental changes such as temperature, affect species and their populations, is central to the process of speciation. In the case of Alvinellidae, previous studies have proven that the long-term adaptation to contrasting thermal regimes has induced specific adaptation at the molecular level, improving the thermostability of proteins in ‘hot’-adapted lineages, whereas other lineages living in colder environments experienced some selective relaxation on their proteomes [15, 17, 77, 95]. Here, for the first time, we resurrected ancestral proteins of two key enzymes to study their thermostability, and demonstrate that alvinellid ancestors were mildly thermophilic. Moreover, we showed that these ancestors underwent parallel adaptive evolution, producing independent lineages of ‘hot’and ‘cold’-adapted species. These nascent species should have lived sympatrically for long enough to exchange alleles, resulting in a mosaic of topological patterns at the genome scale. In addition, by coupling force field simulations with FoldX and inferences on the amino acid usage bias at the proteome level, we confirmed the robustness of the ancestral sequence reconstruction procedure over a large number of genes, regardless of the tested phylogenetic hypothesis. This enabled us to finally reconstruct the adaptive history of proteins in these vent worms spanning more than 100 million years of evolution.

## 6 Supporting Information Appendix

### List of supplementary figures

- Supplementary figure 1 Discriminative topologies used for mapping over the *A. pompejana* genome
- Supplementary figure 2 Alignments statistics: number of transcripts and reciprocal best-hit blast
- Supplementary figure 3 Projected compositions of different classes of amino acid residues for all Terebelloformia species
- Supplementary figure 4.A. p-value distributions of the topologies in 5000 shuffled genomes
- Supplementary figure 4.B. p-value distributions of the gene structure in 5000 shuffled genomes
- Supplementary figure 5 Mapping of gene coalescence onto *A. pompejana* genome
- Supplementary figure 6 Examples of experimental measures for the unfolding of proteins, *P. fijiensis* and *P. palmiformis* cMDH, *P. sulfincola* and *P. pandorae* Cu/Zn SOD
- Supplementary figure 7 Likelihood and negative AIC gain given the number of coordinates
- Supplementary figure 8 Amino acid usage optmized for ancestral species: model optimization
- Supplementary figure 9 Amino acid usage for ancestral species: comparison with standard model
- Supplementary figure 10 Reconstruction of amino acid composition for buried residues, comparison with standard model

### List of supplementary tables

Supplementary table 1 Number of transcripts per species
Supplementary table 2 Number of alignments per discriminative topology
Supplementary table 3 Number of genes used to perform the training of the evolutionary model
Supplementary table 4 Number of positions used to perform the testing of the evolutionary model
Supplementary table 5 SOD thermodynamic parameters
Supplementary table 6 cMDH thermodynamic parameters
Supplementary table 7 R2 regression with Foldx4 with different structure reference
Supplementary table 8 Predicted and measured ΔΔ*G*
Supplementary table 9 AIC gain compared to the standard model for all alignments in all topologies
Supplementary table 10 Coefficients associated with the two thermostablity indices Detailed protocol for the expression, purification and measure of recombinant proteins is also presented as supplementary material.

## 7 Author contributions

PGB: Writing – original draft, Writing – review & editing, Methodology, Formal analysis, Data curation, Software, Investigation. ASP: Investigation. MB: Investigation. SB : Investigation. MAN: Investigation. LC: Investigation. JM: Writing – review & editing, Investigation, Supervision, Funding acquisition. DJ: Writing – review & editing, Supervision, Resources, Project administration, Funding acquisition.

## 8 Declaration of interest

The authors declare that they have no known competing financial interests or personal relationships that could have appeared to influence the work reported in this paper.

## 9 Data availability

Scripts and data used to perform the analysis are available at github.com/Pigui1/Alvinellid ASR

## 10 Acknowledgments

We thank the chief scientists (S. Hourdez, D. Jollivet) and the crew behind the oceanographic cruise Chubacarc (doi:10.17600/18001111), as well as the Roscoff ABIMS platform of bioinformatics. We also thank Murielle Jam (Station biologique de Roscoff) for her precious advice during the purification of the proteins.

## Notes

https://github.com/Pigui1/Alvinellid_ASR

### Competing Interest Statement

The authors have declared no competing interest.

