## Supplementary figures and tables for "Ecological Diversification into Broad Thermal Niches Revealed by Protein Resurrection and Proteome-Wide Ancestral Reconstruction During the Rapid Radiation of Alvinellid Worms"

### Retrieving orthologous genes and *de novo* assembly

We needed to increase the number of transcripts for the species *P. pandorae*, in order to have a greater resolution for the chromosome mapping of gene topologies.

We performed *de novo* assembly using Trinity v. 2.9.1 [1] on the reads of *P. pandorae* from Fontanillas et al. 2017 [2] with a per base phred-score above 25, and the transcripts were filtered with Kraken v.2.0.9 [3] to remove potential contamination. These new transcripts, and the transcripts of *P. pandorae* obtained in Brun et al. 2024 [4] were pooled together with cap3 [5] (35 nucleotide overlap, with 1 mismatch allowed). The transcripts were then predicted with Transdecoder to obtain complete trancripts, or transcripts of more than 200 amino acids.

We also performed a partial assembly of the genome of *P. pandorae* using Illumina sequencing (MiSeq). The reads were assembled with SPADES [6]. This resulted in a very fragmented genome, on which we performed *ab initio* gene prediction using Augustus [7]. Augustus was trained on the first 14000 scaffolds (between 4000 and 35000 nucleotides long) using the complete transcripts longer than 100 amino acids and the corresponding translated proteins first obtained from RNAseq. For the gene prediction, all complete transcripts and transcripts longer than 200 amino acids were used as hints.

Unique predicted genes and transcripts were pooled together, retaining only complete transcripts and transcripts longer than 200 amino acids. The sequences are finally filtered with Kraken to remove any potential contamination by plansmid, virus, fungi, plant, bacteria, archae and *H. sapiens*. This last filtering step is applied to all transcriptome from Brun et al. 2024 [4].

We performed reciprocal best-hit blast against *A. pompejana*, taken as a complete reference of the gene catalogue. The orthologous genes were used to construct small alignments that are able to discriminate between the different toplogies at the root of the Alvinellidae family. All nodes that do not contribute are collapsed, such as we reduce the potential uncertainty that comes from other parts of the trees. We always chose the longest transcript to build the alignments (supplementary table 1).

This strategy allows us to build a high number of discriminative alignments (supplementary figure 1).

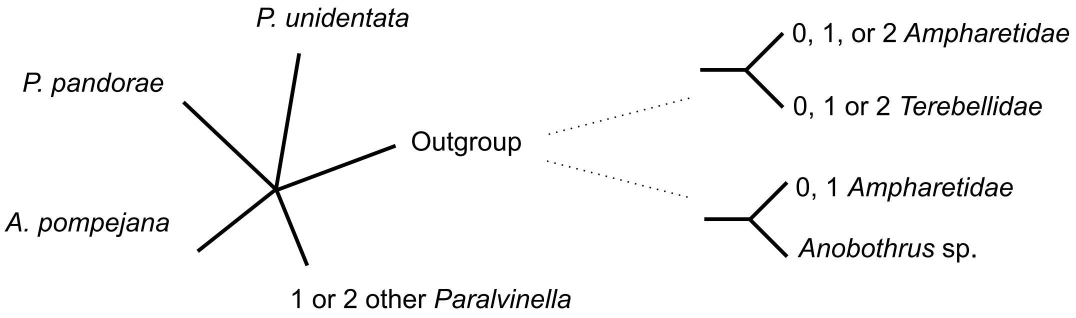


Supplementary figure 1 - Discriminative topologies used to distinguish the phylogenetic hypothesis without added uncertainty.

| | Number of transcripts | Number of reciprocal blast with *A. pompejana* (number of sites) | Number of short blast <150 amino acids (number of sites) |
|---|---|---|---|
| *P. fijiensis* 1 | 77,000 | 10622 (3,782,050) | 1560 (163,308) |
| *P. fijiensis* 2 | 40,971 | 9664 (3,425,619) | 1367 (142,776) |
| *P. fijiensis 3* | 38,873 | 9543 (3,378,196) | 1351 (142,963) |
| *P. fijiensis* 4 | 48,483 | 10071 (3,619,987) | 1431 (148,258) |
| *P. hessleri* 1 | 53,805 | 10556 (3,755,352) | 1476 (155,612) |
| *P. hessleri 2* | 35,894 | 9507 (3,349,081) | 1281 (132,252) |
| *P. mira* | 29,386 | 8515 (2,902,734) | 1190 (133,319) |
| *P. pandorae* | 61,127 | 8497 (2,130,605) | 2982 (275,530) |
| *P. unidentata* 1 | 80,169 | 10983 (3,819,603) | 1644 (169,858) |
| *P. unidentata* 2 | 45,958 | 9786 (3,354,319) | 1429 (146,280) |
| *P. unidentata* 3 | 60,051 | 10276 (3,750,186) | 1461 (150,282) |
| *A. pompejana* | 27,350 | - | - |
| *P. palmiformis* | 46,445 | 10463 (3,826,930) | 1470 (154,134) |
| *P. grasslei* | 22,257 | 7152 (2,422,786) | 992 (105,321) |
| *H. invalida* | 27,484 | 8759 (3,429,978) | 1086 (115,792) |
| *A. gunneri* | 33,233 | 7621 (2,461,623) | 1186 (123,735) |
| *N. edwardsii* | 42,995 | 9191 (3,564,220) | 1208 (129,172) |
| *S. kaia* | 168,229 | 9908 (3,234,156) | 1746 (177,977) |
| *M. palmata* | 90,367 | 9948 (3,803,079) | 1418 (149,815) |
| *P. gouldii* | 11,498 | 2769 (747,382) | 496 (49,344) |
| *A. carldarei* | 19,434 | 5962 (1,744,577) | 1017 (104,901) |
| *P.* sp. nov | 31,494 | 7316 (2,098,543) | 1306 (129.599) |
| *P. sulfincola* | 17,431 | 5405 (1,290,972) | 1430 (140,402) |
| *A. caudata* | 66,690 | 11895 (4,062,569) | 1681 (167,189) |
| *Anobothrus* sp. | 30,709 | 7998 (2,572,816) | 1277 (133,622) |

Supplementary table 1 - Number of transcripts per species

| | Number of orthologous genes | Number of sites | number of orthologous genes > 150 amino acids or complete | Number of sites in alignments longer than 150 amino acids or complete |
|---|---|---|---|---|
| 4 Alvinellidae + 1 Outgroup | 228 | 40,688 | 147 | 35,813 |
| 5 Alvinellidae + 1 Outgroup | 259 | 51,015 | 194 | 50,359 |
| 4 Alvinellidae + several Outgroup | 450 | 87,458 | 325 | 82,148 |
| 5 Alvinellidae + several Outgroup | 4,884 | 1,020,845 | 3, 448 | 912,125 |
| Total | 5,821 | 1,200,006 | 4,114 | 1,080,445 |

Supplementary table 2 - Number of alignments per discriminative topology

We also added 433 orthologous groups of genes from Brun et al. 2024 [4], identified with Orthograph [8] instead of reciprocal blast, and not overlapping with the new orthogroups built for this study. In total, we obtained 6254 genes (and 385 other genes without outgroup species, which have a low discriminative power, supplementary table 2).

The likelihood of the different topologies are tested one against each other on discriminative topologies with IQ-Tree (model LG+G+F) [9].

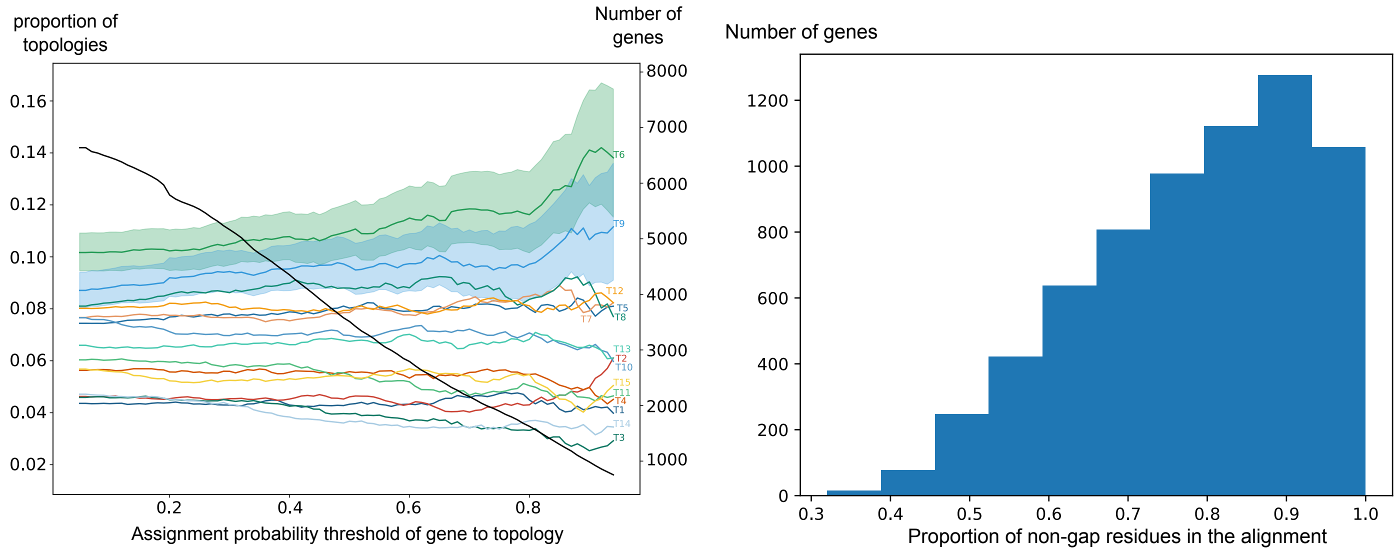


Supplementary figure 2 - Alignments statistics. Left panel: expected fraction of genes belonging to each topology. A filter is applied to remove genes that are weakly associated with a topology, from a minimum probability of 0.05 to 0.95. The total number of genes thus varies from 6639 to 753. 95% Wilson confidence intervals are displayed for topologies T6 and T9. Right panel: distribution of the genes given the completude of the alignment. Most alignments have above 80% completness.

**Training and testing of 15 evolutionary models with amino acid variation**

Finally, we gather the genes given their highest posterior probability to belong to each topology (supplementary table 3). Genes with a probability smaller than 0.5 are excluded. We retrieved the full alignments with extended topologies (allowing a maximum of two missing species), leaving 2544 orthologous groups. These groups of genes were concatenated and used to train the phylogenetic models with amino acid frequencies variation.

| | Number of genes | Alignment length (amino acids) |
|---|---|---|
| T1 | 115 | 8866 |
| T2 | 95 | 6554 |
| T3 | 107 | 7746 |
| T4 | 135 | 11091 |
| T5 | 208 | 14552 |
| T6 | 282 | 21561 |
| T7 | 214 | 20399 |
| T8 | 238 | 25888 |
| T9 | 265 | 26845 |
| T10 | 178 | 12879 |
| T11 | 132 | 15621 |
| T12 | 203 | 14452 |
| T13 | 165 | 14947 |
| T14 | 97 | 7915 |
| T15 | 110 | 7262 |
| Total | 2544 | 216578 |

Supplementary table 3 - Number of genes used to perform the training of the evolutionary model with amino acid variation.

The models are later tested on all 2011 non informative genes, for which the posterior probability to be associated with a particular topology is below 0.5. Only 263 of these genes contain all species (400 with 1 species missing, 331 with two species missing). The 263 genes with complete species are concatenated together, and we only kept positions in which every species are present to build alignments dividing the positions according to the kind of structure the belong to (supplementary table 4).

| | | Alignment length (amino acids) |
|---|---|---|
| Exposed | Helix | 5246 |
| | Sheet | 1189 |
| | Coil | 8355 |
| Buried | Helix | 4029 |
| | Sheet | 2657 |
| | Coil | 2032 |
| Total | | 23508 |

Supplementary table 4 - Number of positions used to perform the testing of the evolutionary model with amino acid variation.

The different alignments are plotted against two axis expected to correlate with the stability of the proteins (supplementary figure 3).

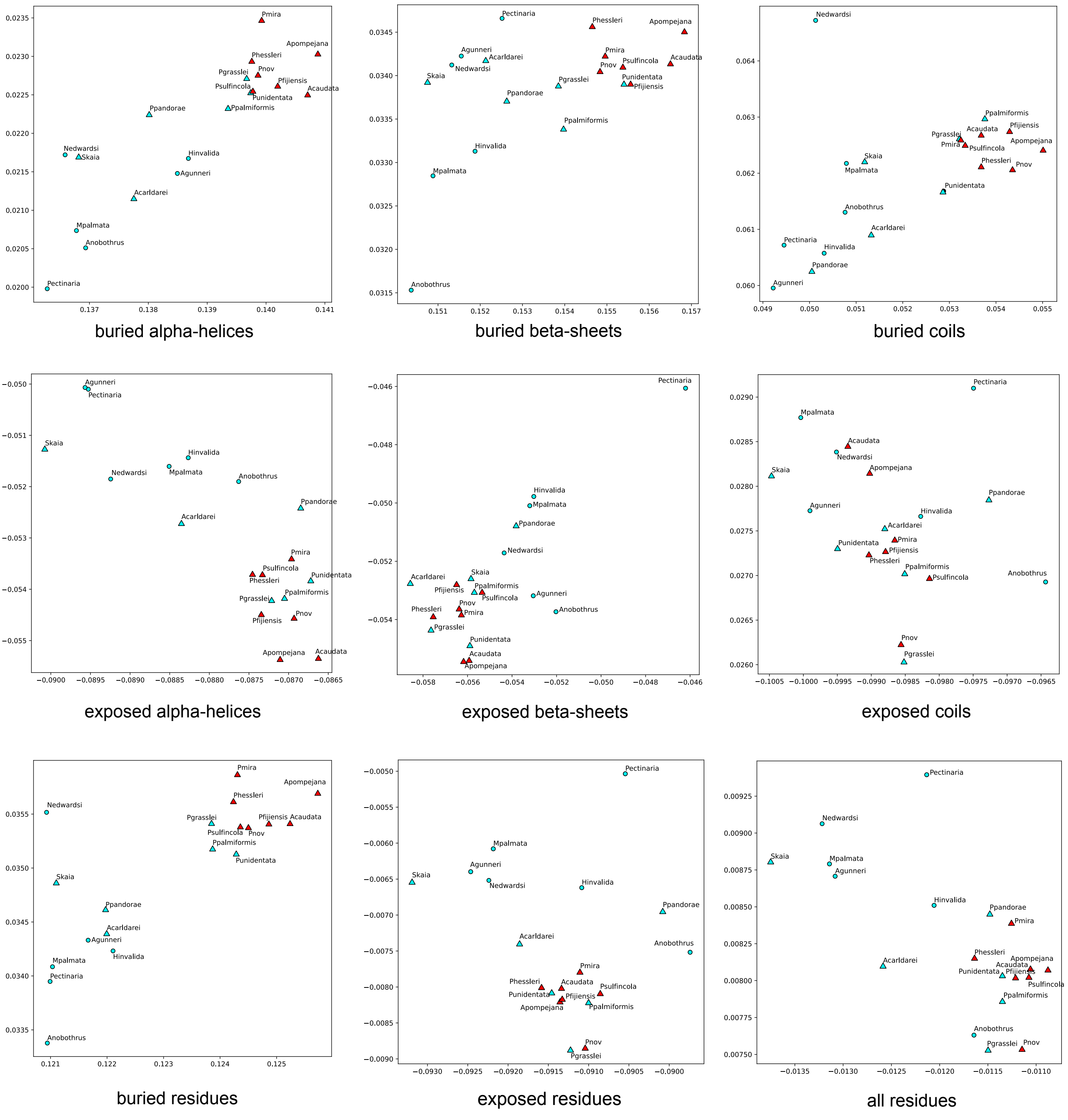


Supplementary figure 3 - Projected compositions of different classes of amino acid residues in Terebelloformia species, according to the first principal components identified in Tsuboyama et al., Nature, 2023 [10] figure 3. Hydrothermal species have a triangle, warm and cold species are in red and blue.

### Mapping of the genes on the genome of *A. pompejana*

The test for over-representation of genes along the genomes are described in supplementary figure 4, while the full mapping of gene topologies over the genome of *A. pompejana* [11] is given in supplementary figure 5.

The unequal distribution of topologies ($T$) along the genome was tested using binomial tests ($\mathscr{B}_T(n, p)$), where $n$ is the number of genes found in a sliding window of 2.5 Mb along the chromosomes (1231 windows covering the entire genome with 48.5 genes on average), and $p$ is the empirical frequency of a given topology $T$ over all genes. The null hypothesis is that the observed distribution of $T$ over all genomic windows falls within the expected fluctuations of $T$ frequencies. To do so, we tested if any topology is over-represented in one or more genomic windows. The required p-values to reject the null hypothesis were determined with 5000 simulations, in which the gene topologies and the gene positions were shuffled. The lowest $p_T$ p-values were collected independently for each topology over all genomic windows in each simulation. $p_T$ was exponentially distributed, and this distribution was used to approximate the required threshold to reject the null hypothesis and conclude that $T$ is over-represented in a given window, with a less than 5\% chance of any topology to be falsely positively identified on any window of the genome.

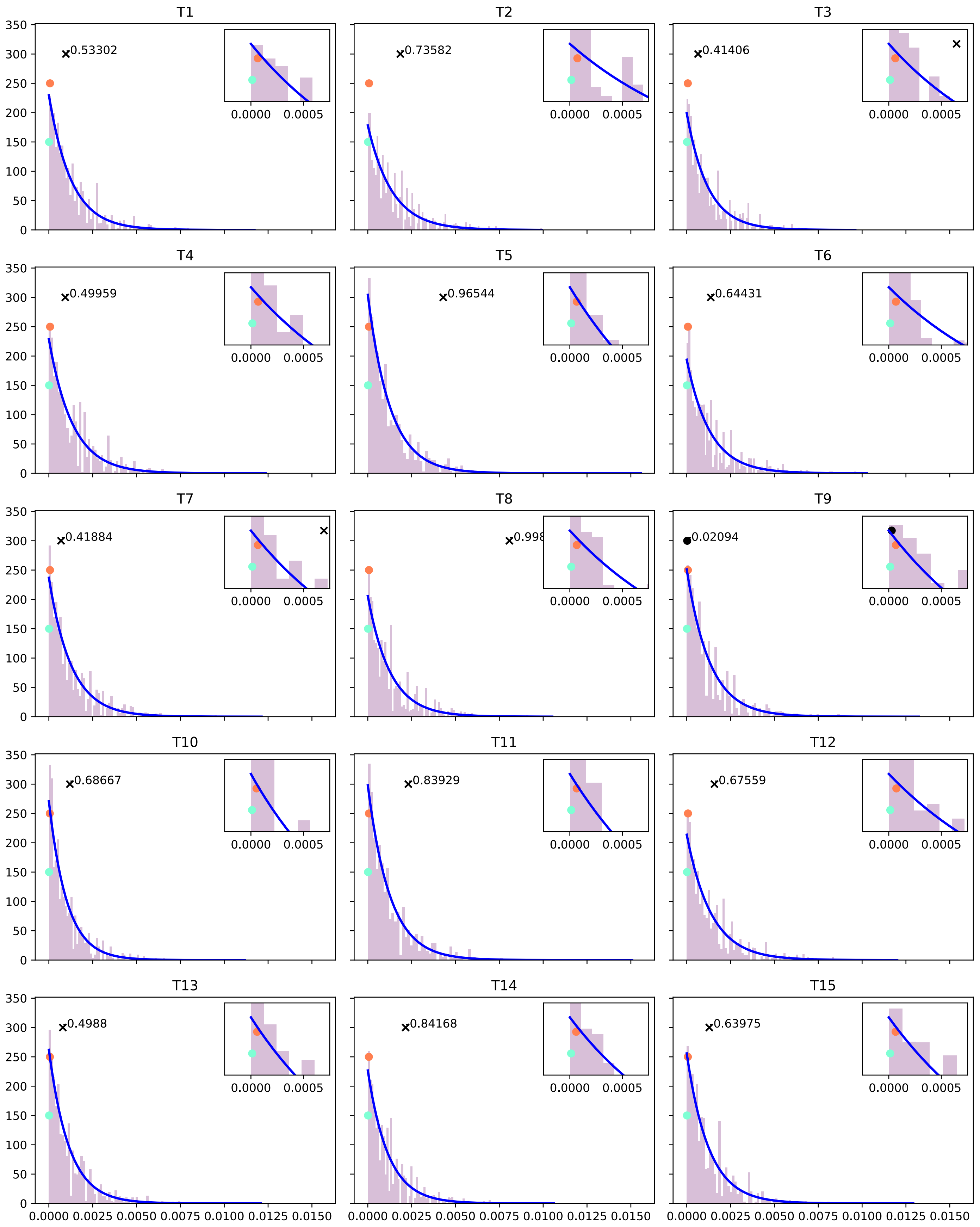


Supplementary figure 4.A - p-value distributions of the topologies in 5000 shuffled genomes. For each shuffled genome, the lowest p-value reached by a topology on all genomic windows is retained. The 5000 p-values per topology are plotted in the purple histogram, and give the distribution of the null hypothesis that the high number of genes from a specific topology in a genomic window is only due to chance. Green dots correspond to the lowest 1% of p-values, Red dots the lowest 5.0% from a fitted exponential law. Black dots are the actual p-value met by one toplogy on the original, not shuffled genome. Dots indicate that the p-value is significant at a 5% threshold, cross that it does not meet the criteria.

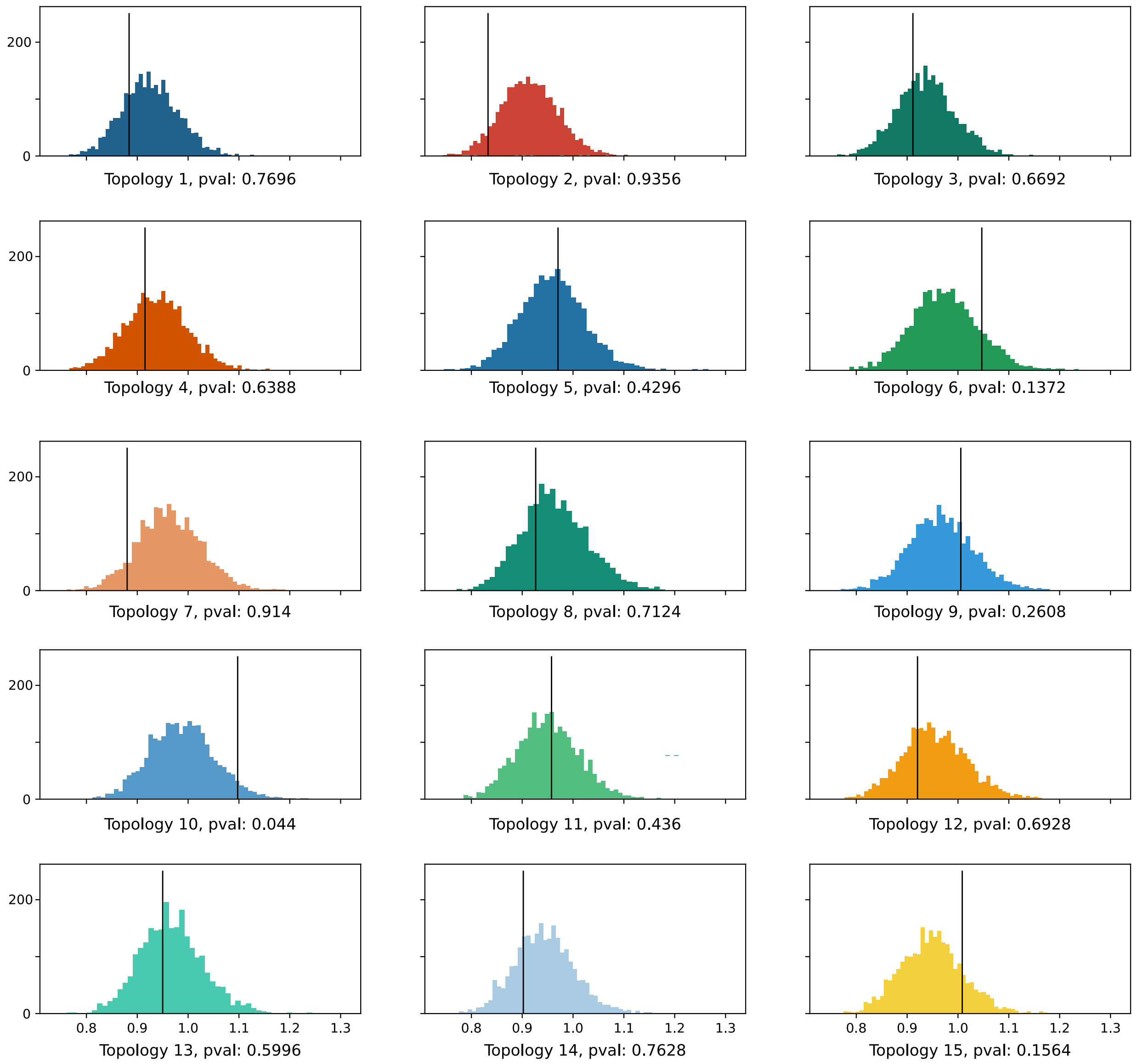


Supplementary figure 4.B - p-value distributions of the gene structure in 5000 shuffled genomes. For each shuffled genome, we gather the distribution of the deviation to the expected average of the oberved number of topologies across all genomic windows. These distributions are fitted to exponentials, which can be solely characterized by their means. A higher mean implies a more structured genome, with a higher number of genomic windows with a very high or very low occurence of a given topology. The distribution of means for all shuffled genomes is shown, as well as the empirical mean of the true genome as a black line. A more-than-expected structured empirical genome would imply a higher mean than random simulations.

The 5000 simulations were also used to construct a second test, which does not aim at identifying a precise topology imbalance within a given genomic window, but instead characterizes the global imbalance of a given topology over the whole genome. The null hypothesis is that $T$ is equally interspaced over the genome. This test aggregates weaker statistical information at the expense of resolution. In that case, the $p$ values of a given topology are collected over all genomic windows for each simulation. $p$ is remapped such as if $p > 0.5, p \leftarrow -p + 0.5$, so that $p$ is close to $0$ if the topology is over or underrepresented on the genomic window, and $0.5$ if it is at the expected average frequency. $x = |log(p)| + log(0.5)$ is well approximated by an exponential distribution $\lambda exp(-\lambda x)$, which is entirely characterized by its mean $m = 1/\lambda$. A lower mean indicates more windows with a probability closer to $0.5$, thus close to the expected average, while a higher mean indicates a deviation toward more unbalanced genomic windows. The 5000 simulations account for a distribution of 5000 simulated means, against which we can compare the observed mean of the true data. If the true data has a high mean compared to shuffled genomes, then the localization of the topology is less homogeneous than expected by random.

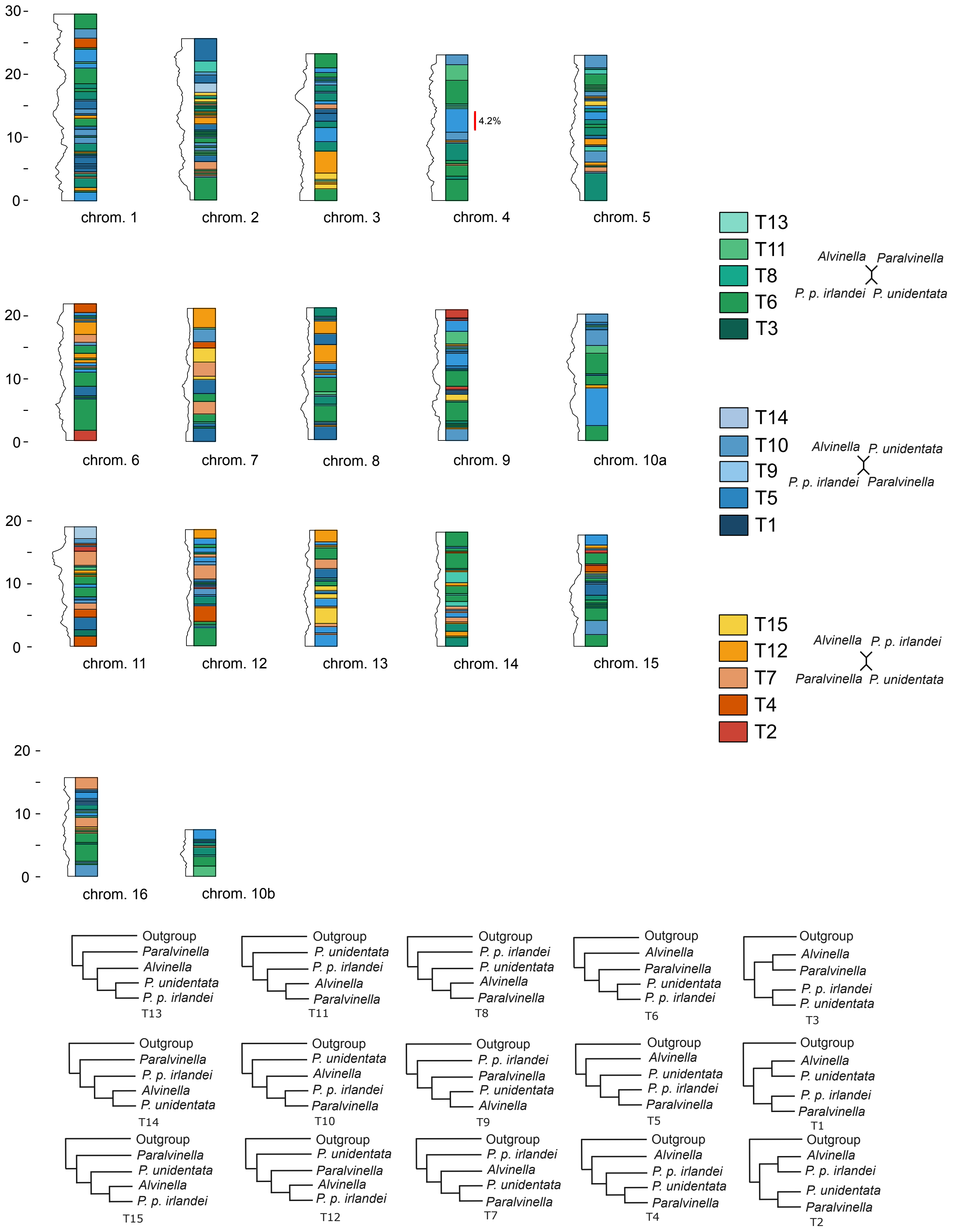


Supplementary figure 5 - Mapping of gene coalescence onto *A. pompejana* genome. The most abundant gene topology on a window of 2.5Mb is displayed, sliding every 0.25Mb. The chromosom length is given in Mb. The relative number of aligned nucleotides on each sliding window is given with the black enveloppe (peaking at 57.749 residues on chromosom 11). Portions of the chromosoms were the T9 topology is over-represented is given on the rigth side. Topologies are gathered in green/blue/red according to the simplest topology between the 4 alivinnelid lineages.

**Expression of recombinant proteins**

cMDH and Cu/Zn SOD sequences of a series of 12 modern alvinellid species and ancestral sequences were reconstructed from the species phylogenetic tree using different ASR algorithms.

All modern and ancestral sequences were produced as *E. coli* recombinants in a heterologous system. The expressed contemporary species were the thermophilic species *A. pompejana, P. sulfincola, P. fijiensis, P. mira*, and the cold-adapted species *P. grasslei, P. palmiformis, P. p. irlandei* and *P. unidentata*. The expressed proteins for ancestors corresponded to all internal nodes of the Alvinellidae tree, excluding the last common ancestor (LCA) of *A. pompejana* and *A. caudata*, the LCA of *P. palmiformis* and *P. grasslei*, the LCA of *P. hessleri* and *P. mira*, and the LCA of *P.* sp. nov. and *P. fijiensis*, as these species pairs are very close to one another and share similar thermal habitat. Both ancestral sequences reconstructed under either the T6 or T9 hypotheses were considered for the protein expression. For each protein construction, T7 Express lysY/Iq Competent *E. coli* (NEB C3013) were transformed with pET100/D-TOPO expression vectors (ThermoFisher) containing the protein coding sequence fused with a (His)6 tag. The bacteria were grown in a LB medium containing 100 µg/mL of ampicillin and the expression was induced by injecting 1 mM IPTG for 4 hours at 37°C until the OD600 reached between 0.4 and 0.6. Cultures were then harvested, and pelleted cells were re-suspended into a His-Bind Buffer of the Novagen™ His-Bind Kit kit for the purification steps. Bacteria were sonicated and a 1‰ DNAse A was added for 30 min on ice. The lysate was centrifuged and the proteins were purified from the supernatant using a $Ni^{2+}$ chelation chromatography on 2 mL columns with the His-Bind Kit of Novagen following an elution using a 20 mM Tris-HCl, 500 mM NaCl, 500 mM Imidazole pH 7.4 buffer. A second purification step was performed by a size-exclusion chromatography with a HiLoad 16/60 Superdex 75 (Cytiva) column connected to an ÄKTA Avant system in 20 mM Tris-HCl, 200 mM NaCl pH 7.4 buffer.

The elution was monitored at 280 nm for cMDH and 205 nm for the SOD. Proteins were then concentrated between 0.6 and 1.4 mg/mL, depending on the total quantity of harvested recombinant bacteria. Half-denaturation temperatures of the cMDH proteins were measured by nano-format of Differential Scanning Fluorimetry (nanoDSF) using the Prometheus NT48 (Nanotemper). This technique requires low amounts of proteins (30 µg for one experiment in triplicate), and is well suited for the cMDH which possess 6 tryptophane residues. The 350/330 nm fluorescence ratio, $r(T)$ (unveiling of the W residues) was measured from 20°C to 95°C with a temperature increase of 1°C/min.

$r(T)$ is linearly correlated with the fraction of unfolded protein $f_u(T)$ with the correction established by Yadav et Ahmad (2000) [12]:

$$r(T) = f_u(T) \times (a_u \times T + b_u) + (1 - f_u(T)) \times (a_n \times T + b_n)$$

Where $a_u$, $b_u$, $a_n$, and $b_n$ are scalars adjusting for the linear correction of the signal.

The fraction of unfolded proteins was obtained from the equilibrium constant $K_{eq}(T)$ according to:

$$f_u(T) = \frac{K_{eq}(T)}{1 + K_{eq}(T)}$$

The equilibrium constant $K_{eq}(T)$ is linked to the standard free energy variation during unfolding (denaturation) step $\Delta G^o(T)$ by:

$$K_{eq}(T) = exp(-\frac{\Delta G^o(T)}{RT})$$

$R$ is the molar gas constant. Finally, $\Delta G^o(T)$ was obtained with the Gibbs-Helmholtz relationship [13]:

$$\Delta G^o(T) = \Delta H_m \times [1 - \frac{T}{T_m}] - \Delta C_p \times [T_m - T + T \times ln(\frac{T}{T_m})]$$

where $T_m$ is the melting temperature at which half of the proteins are denatured, $\Delta H_m$ the enthalpy variation at $T_m$, and $\Delta C_p$ the heat capacity change of the protein during unfolding. Combining these relations allows to fit $f_u(T)$ against $T$ and determine the parameters during the *in vitro* measurements of the protein denaturation:

$$f_u(T) = \left[1 + exp\left(\frac{\Delta H_m}{R}\left(\frac{1}{T} - \frac{1}{T_m}\right) - \frac{\Delta C_p}{R}\left(\frac{T_m}{T} - 1 + \ln\frac{T}{T_m}\right)\right)\right]^{-1}$$

Thermodynamic parameters for the Cu/Zn SOD were measured by differential scanning micro-calorimetry using the Microcal PEAQ-DSC (Malvern), as these proteins do not contain tryptophanyl residue to monitor its fluorescence with nanoDSF. The excess heat capacity $C_p^{exp}$ was measured between 20-110°C, with a temperature increase of 1°C/min.

In this case, $C_p^{exp}(T)$ is related to the fraction of unfolded protein by [14]:

$$C_p^{exp}(T) = \Delta H_c \frac{df_u}{dT}$$

with $\Delta H_c$ the calorimetric enthalpy variation. Using the relation between $f_u$ and $K_{eq}$:

$$\frac{df_u}{dT} = \frac{1}{(1 + K_{eq}(T))^2} \times \frac{dK_{eq}}{dT}$$

And

$$\frac{dK_{eq}}{dT} = K_{eq}(T) \times (\frac{\Delta G^o(T)}{RT^2} - \frac{d\Delta G^o}{RTdT})$$

$\Delta G^o(T)$ is obtained from the Gibbs-Helmholtz relation, and consequently

$$\frac{\Delta G^o}{dT} = \frac{\Delta H_m}{T_m} - \Delta C_p \times ln(\frac{T}{T_m})$$

These equations can be used to express $C_p^{exp}(T)$ against $T$, depending on $\Delta H_c$, $T_m$, $\Delta H_m$ and $\Delta C_p$, which can be determined from the experimental curves:

$$C_p^{exp}(T) = \Delta H_c f_u(T)[1 - f_u(T)]\left[\frac{\Delta H_m}{RT^2} - \frac{\Delta C_p}{RT}(1 - \frac{T}{T_m})\right]$$

We considered three potential successive denaturation events for the SOD [15]. The measured signal is consequently assumed to be the sum of three distinct $C_p^{exp}$, with their own thermodynamic parameters to estimate.
Denatruation curves and thermodynamic parameters are given in supplementary figure 5 and supplementary tables 5 and 6).

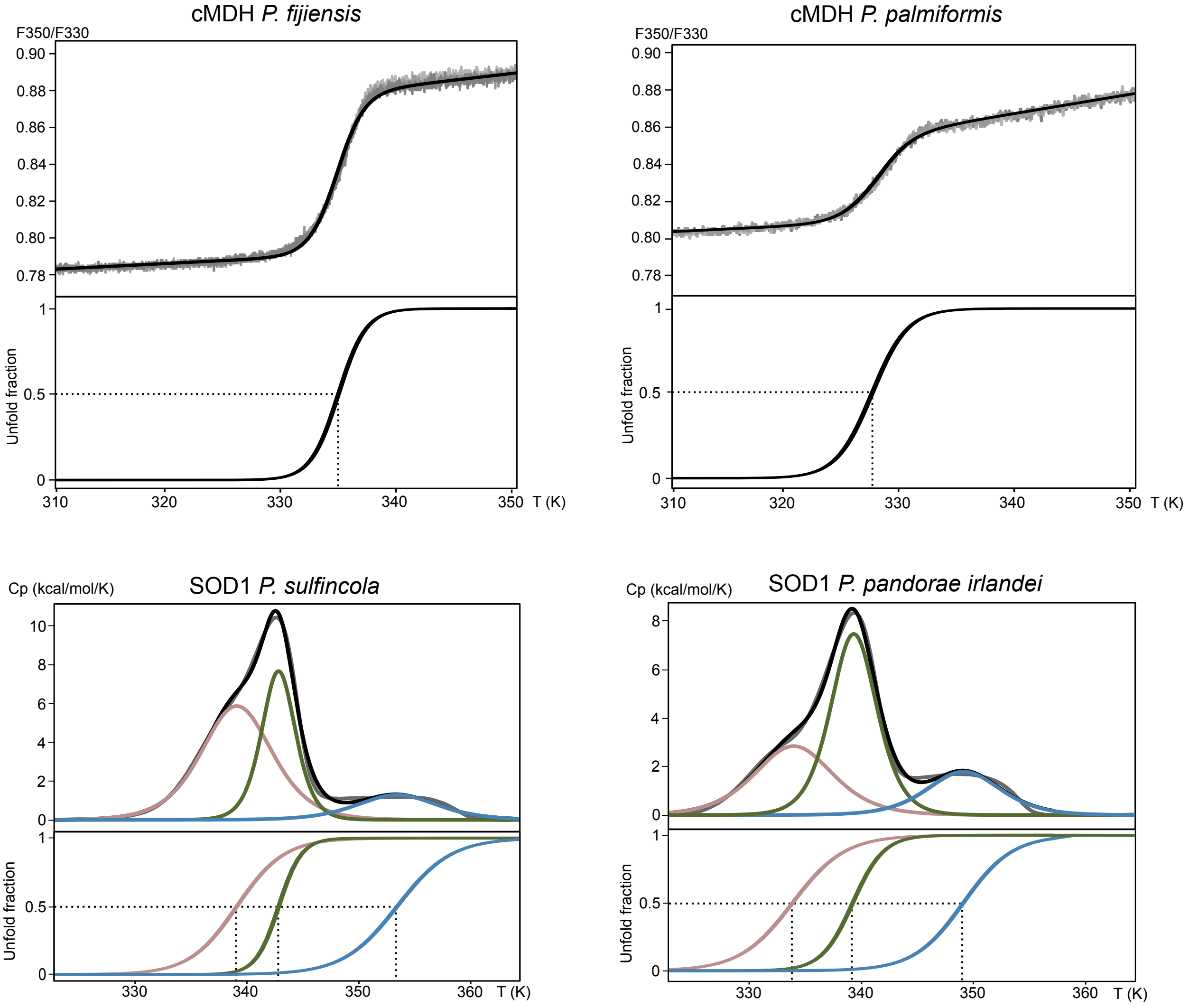


Supplementary figure 6 - Experimental measures for the unfolding of proteins. Top panels: Denaturation curves obtained by nanoDSF for two cMDH proteins of *P. fijiensis* and *P. palmiformis* (350/330 nm fluorescence ratio against temperature). Grey lines : experimental curves. Black line : theoretical fit. Bottom panels show the integral of the theoretical denaturation curve, with the Tm indicated by dotted lines. Bottom panels: Denaturation curves obtained by nanoDSC for SOD proteins of *P. sulfincola* and *P. p. irlandei* (heat capacity against temperature). Grey line : experimental curve. Black line : theoretical fit. Red, green and blue lines : decomposition of the theoretical fit into three denaturation events. Bottom panels show the integral of the theoretical denaturation curves, with the Tm indicated by dotted lines.

| | ΔHm (kJ/mol) | ΔCp (kJ/mol/K) | $ΔCp^{adj}$ (kJ/mol/K) | Tm (°C) |
|---|---|---|---|---|
| *P. unidentata* | 739 | 31 | 24 | 59 |
| *P. pandoae* | 666 | 26 | 21 | 61 |
| *P. grasslei* | 678 | 29 | 22 | 57 |
| *P. palmiformis* | 648 | 28 | 22 | 56 |
| *A. pompejana* | 753 | 26 | 21 | 66 |
| *P. mira* | 719 | 29 | 22 | 61 |
| *P. fijiensis* | 747 | 28 | 23 | 62 |
| *P. sulfincola*/Anc5-T6/Anc5-T9 | 702 | 26 | 21 | 62 |
| Anc1-T6 | 738 | 29 | 22 | 62 |
| Anc2-T6/Anc2-T9/Anc6-T9 | 757 | 30 | 24 | 60 |
| Anc3-T6/Anc3-T9 | 672 | 26 | 21 | 61 |
| Anc4-T6/Anc4-T9 | 717 | 27 | 22 | 62 |
| Anc6-T6 | 682 | 27 | 21 | 60 |
| Anc1-T9 | 772 | 31 | 24 | 60 |

Supplementary table 5 - cMDH thermodynamic parameters

| | ΔHm (kJ/mol) | ΔCp (kJ/mol/K) | $ΔCp^{adj}$ (kJ/mol/K) | Tm (°C) |
|---|---|---|---|---|
| *P. unidentata* | 481 | 17 | 12 | 76 |
| *P. pandorae* | 423 | 17 | 10 | 75 |
| *P. grasslei* | 642 | 0 | 0 | 75 |
| *P. palmiformis* | - | - | - | 72 |
| *A. pompejana* | 481 | 17 | 11 | 80 |
| *P. fijiensis* | 442 | 19 | 10 | 82 |
| *P. sulfincola* | 386 | 15 | 9 | 80 |
| *P. mira* | - | - | - | - |
| Anc1-T6/Anc2-T6 | 488 | 21 | 11 | 79 |
| Anc3-T6/Anc4-T6 | 464 | 22 | 11 | 79 |
| Anc5-T6/Anc5-T9 | 295 | 10 | 7 | 83 |
| Anc6-T6 | 609 | 32 | 14 | 79 |
| Anc1-T9/Anc2-T9 | 528 | 26 | 12 | 78 |
| Anc3-T9/Anc4-T9 | 464 | 22 | 11 | 79 |
| Anc6-T9 | 387 | 17 | 9 | 80 |

Supplementary table 6 - SOD thermodynamic parameters corresponding to the 3rd denaturation event (blue line on supplementary figure 6)

### Simulation of ancestral proteins

We compared the experimental measured stabilities and the Foldx4 [16] predicted stabilities, which should be linearly correlated. Foldx4 needs to use one protein 3D template corresponding to a reference sequence in order to predict the effect of point mutations on the stability. To this end, we predicted the structure with Colabfold [17] of several sequences for the two protein families, for which we expect the structure to be « in the middle of the pack », namely the two most ancestral sequences Anc1 under the two phylogenetic hypothesis, and a consensus sequence from all modern and ancestral ML sequences (suppplementary table 7). We chose to allow 1 potential outlier in each regression, that is excluded, we kept the protein structure that had the best prediction power given the potential variery of infered sequences.

The predictions are best aligned with measures at simulated temperature of 48.5°C for the cMDH, and 82.7°C for the SOD, according to the Gibbs-Helmots equation

$$\Delta G^o(T) = \Delta H_m \times (1 - T/T_m) - \Delta C_p \times (T_m - T + T \times ln(T/T_m))$$

The predictions are therefore in good agreement with experimental measures around the $T_m$ of the two protein families respectively. This hints that our measures of $\Delta G^o$ are of good quality around 0 but diverge quickly, likely because the fitting of the curves is robust for $T_m$ and $\Delta H_m$, but we lack precision for $\Delta C_p$. For this reason, we later consider an adjusted $\Delta C_p$ to avoid a cold denaturation above 0°C, according to [18].

The predictions on modern species shows three distinct groups of species with increasing protein stability: non-alvinellid and colder species, colder alvinellid species and warm alvinellid species (supplementary table 8). A few species have surprising predictions: SOD *P. mira* and *P. hessleri*, predicted as colder, and MDH *P. fijiensis* and *P*. sp. nov. Only one measure is available for the P. fijfiensis MDH, which disagrees with the prediction and confirms that this protein is very stable. *P. mira* and *P. hessleri* are sister-species in the phylogeny, and so are *P. fijiensis* and *P*. sp. nov. It is likely that the predictions for these proteins are wrong, because of shared mutations shared by the proteins that FoldX is unable to evaluate correctly.

| | | **Consensus** | **Anc1-T6** | **Anc1-T9** |
|---|---|---|---|---|
| **cMDH** | Repaired PDB, ΔCp adjusted | 0.47/0.79/0.91 | 0.55/0.76/0.88 | 0.53/0.93/0.95 |
| **Cu/Zn SOD** | Repaired PDB, ΔCp adjusted | 0.73/0.80/0.87 | 0.70/0.84/0.87 | 0.69/0.79/0.85 |

Supplementary table 7 - $R^2$ regression with Foldx4 with different structure reference for SOD and cMDH, with all proteins/allowing 1 outlier/allowing 2 outliers.

| | SOD (ΔΔG kcal/mol) | | MDH (ΔΔG kcal/mol) | | |
|---|---|---|---|---|---|
| | Foldx | nanoDSF | Foldx | nanoDSC | |
| *A. pompejana* | 0.0 | 0.0 | 0.0 | 0.0 | 'Hot'-adapted Alvinellidae |
| *A. caudata* | 0.0 | | -0.1 | | |
| *P. fijiensis* | -0.2 | -0.6 | 4.2 | 1.3 | |
| *P. sulfincola* | -0.2 | 0.0 | 2.0 | 1.6 | |
| *P. mira* | 2.0 | | 1.5 | 1.9 | |
| *P. hessleri* | 2.0 | | 1.4 | | |
| *P.* sp. nov. | -1.1 | | 4.2 | | |
| *P. grasslei* | 1.5 | 2.5 | 3.6 | 3.3 | 'Cold'-adapted Alvinellidae |
| *P. palmiformis* | 1.0 | | 3.8 | 3.8 | |
| *P. unidentata* | 1.5 | 1.5 | 2.6 | 2.5 | |
| *P. pandorae* | 1.9 | 1.5 | 2.5 | 2.2 | |
| *H. invalida* | 3.5 | | 10.6 | | 'Cold'-adapted Outgroup Terebelliformia |
| *A. gunneri* | 4.7 | | 8.8 | | |
| *Anobothrus* sp. | 6.9 | | 10.6 | | |
| *A. carldarei* | 3.7 | | 12.8 | | |
| *N. edwardsii* | 4.0 | | 9.1 | | |
| *M. palmata** | 5.1 | | 10.3 | | |
| *S. kaia** | 5.4 | | 6.7 | | |
| *P. gouldii** | 9.1 | | 15.8 | | |

Supplementary table 8 - Predicted (after linear regression) and measured ΔΔG, taking *A. pompejana* as a reference. For some terebellid species marked with (*), a few adustments were made in the SOD WT sequence to adjust the aligned sequence on the 3D template used by FoldX. Species expected to be thermotolerant are in red, colder hydrothermal species in green, other colder species in blue.

### Evolutionary model with amino acid variation

<u>Protocol</u>

Ancestral residue states in a given sequence are reconstructed using a custom-made generalised ancestral sequence reconstruction method that considers the evolution of amino acid composition over time.

It is standard in phylogenetics and ASR to consider the mean usage of amino acid across observed species as a good proxy for ancestral sequences as well. In that case, a standardized mutation matrix, such as the JTT, WAG or LG matrix, is shifted with the frequencies of amino acids:

$$Q' = \pi_{aa} \times Q$$

$Q$ is the standard mutation matrix, $\pi_{aa}$ the empirical frequencies of amino acids, $Q'$ the matrix describing the mutation probabilities in the model. That way, over infinite times, the average composition of the sequences will equilibrate with the average composition of observed species.

Our algorithm leverages this hypothesis by introducing new parameters that allow the equilibrium frequencies to vary over time. To this end, we consider the fluctuation of amino acid equilibrium frequencies in modern species. The bias of amino acid usage is decomposed by principal components analysis (PCA) on all genes where no missing species is allowed at any position of the multiple sequence alignment. The eigenvectors corresponding to the PCA allow to write the amino acid usage of any species according to:

$\overrightarrow{\pi_s} = \overrightarrow{m} + \overrightarrow{v} \cdot \overrightarrow{c_s}$

$\overrightarrow{\pi_s}$ is the amino acid usage of a species, $\overrightarrow{m}$ the mean amino acid usage over the alignment, $\overrightarrow{v}$ the vector base of the PCA, $\overrightarrow{c_s}$ the coordinates of the species in the base.

The 19 coordinates of each ancestral species, corresponding to the 19 degrees of freedom of amino acid usage, in the PCA space can be optimized by maximum likelihood. This adds 19 parameters on each branch, as well as 19 parameters at the root of the tree.

In practice, we reduce the number of parameters by approximating $\overrightarrow{\pi_s}$ with only the coordinates corresponding to the largest eigenvectors. Very small approximated amino acid frequencies which can arise from a linear decomposition during optimization, and they are given a pseudo-count of 0.01% to avoid numerical issues.

Transition probabilities are thus obtained with $Pr_{des} = Pr_{anc} \cdot exp(t \times \pi_s \times Q)$

$t$ is the branch length, $Q$ a standard mutation matrix that we chose to be LG.

The number of coordinates to keep on each branch is chosen iteratively according to the Akaike Information Criterion, which weights the likelihood gain of the model against the number of added parameters:

$AIC = 2 \times (k - L)$, with $k$ the number of parameters and $L$ the optimized likelihood of the model. A model minimizing this quantity is the most informative.

Finally, we also add the parameter $\Gamma$, which describes a probability distribution of the conservation of sites along the alignment. The distribution is dicretized in 4 classes, and the shape of the distribution is optimized by maximum likelihood:

$Pr_{des} = 0.25 \times Pr_{anc} \cdot \sum_{i=1}^{4} exp(g_i \times t \times \pi_s \times Q)$, $g_i$ is the mean of the $\Gamma$ distribution in class $i$.

Models are optimized, and the marginal reconstruction of sequences are taken to compare the evoltuion of amino acid frequencies between modern and ancestral species.

Examples of results fully optimized under the T9 phylogeny (number of axis and eigenvectors) are given below, in supplementary figure 6, 7 and 8.

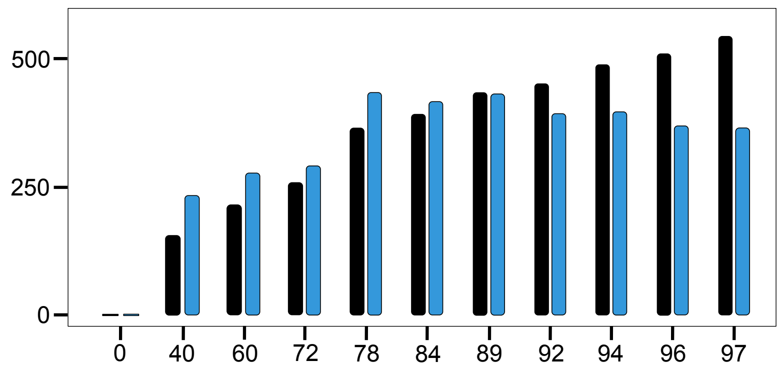


Supplementary figure 7 - Likelihood gain (black) and negative AIC gain (blue) given the number of coordinates added in the model. The cumulative explained variance of the amino acid usage between modern species is given under the axes. The model reaches a optimum at 4 axes (78% of total variance)

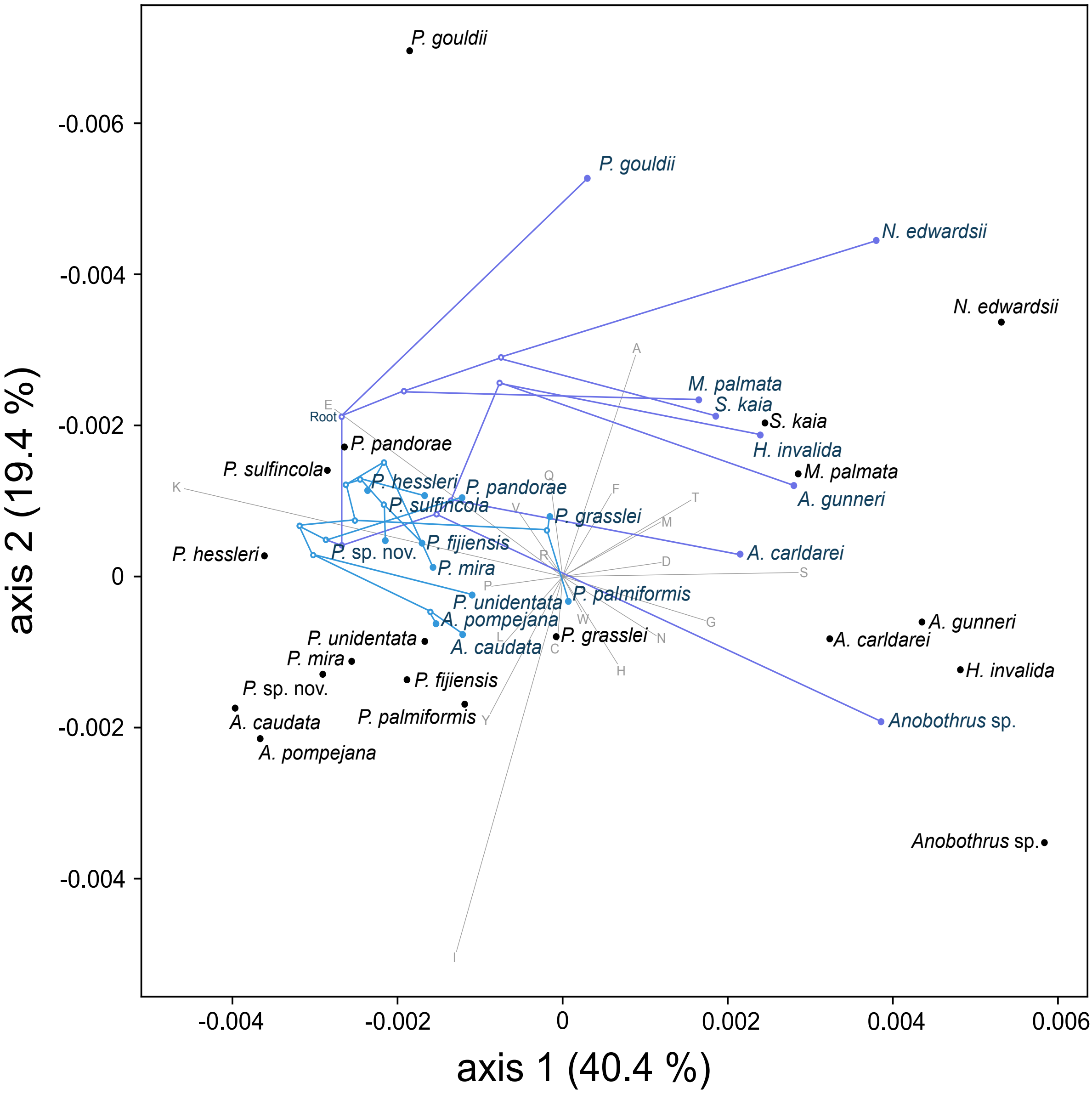


Supplementary figure 8 - PCA of the amino acid usage in modern and ancestral species, optmized with amino acid variations and taking the topology T9 as a reference and 4 coordinates per species. In a standard evolutionary model, all ancestral and modern species would be in the coordinates (0,0), as we consider that all compositions are homogenous, equal to the mean frequencies on the alignment.

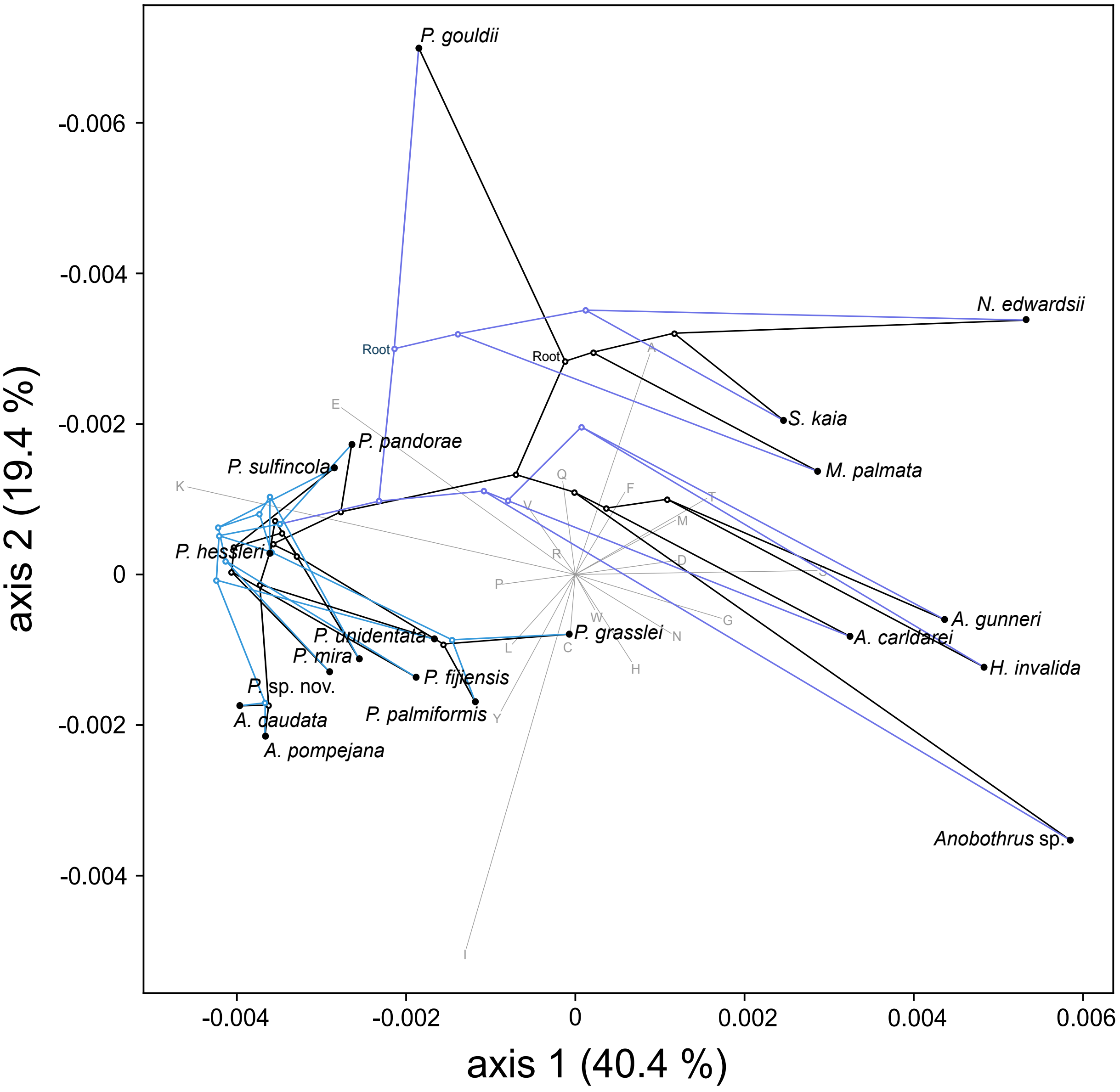


Supplementary figure 9 - Amino acid usage for ancestral species and projected in a PCA space, taking the topology T9 as a reference and 4 coordinates per species. The compositions of modern species are anchored on the input alignment, and the ancestral compositions is inherited from this additional constraint. Black: reconstruction without additional modeling of the amino acid variation. Blue: reconstruction with added variation.

Optimisation on different sequence alignments

The model is optimized independantly on the alignments of protein in agreement with each of the15 topologies. We add PCA axis until the AIC criterion decreases, at which point we stop trying more models. The results of these optimization, compared to a standard model with 0 axis, is given in supplementary table 9. To simplify, we have retained models with 2 axes, except for T2, T3, T14 and T15 where we keep the standard models because of the very small improvement of likelihood added by the new parameters.

| | 0 axis | 1 axis | 2 axes | 3 axes | 4 axes |
|---|---|---|---|---|---|
| # parameters | 36 | 72 | 108 | 144 | 180 |
| T1 | 0 | 23 | 37 | 33 | 3 |
| T2 | 0 | 10 | -34 | -55 | - |
| T3 | 0 | 19 | 5 | -24 | - |
| T4 | 0 | 151 | 145 | 105 | - |
| T5 | 0 | 79 | 51 | 26 | - |
| T6 | 0 | 136 | 132 | 119 | - |
| T7 | 0 | 214 | 240 | 198 | 174 |
| T8 | 0 | 255 | 254 | 223 | - |
| T9 | 0 | 185 | 181 | 168 | - |
| T10 | 0 | 161 | 159 | 142 | - |
| T11 | 0 | 259 | 254 | 226 | - |
| T12 | 0 | 67 | 64 | 32 | - |
| T13 | 0 | 141 | 123 | 77 | - |
| T14 | 0 | 20 | 9 | -25 | - |
| T15 | 0 | -1 | -3 | - | - |

Supplementary table 9 - AIC gain compared to the standard model for all alignments in all topologies, in a common PCA space obtained on the overall alignment. We test an increasing number of axes, until we don't see any improvement in the models anymore. By matter of simplicity, we chose to keep all models with 2 axes to perform ancestral reconstruction, which are amongst the best models, except for topologies T2, T3, T14 and T15 in which we keep the standard model as adding the amino acid variation improves very weakly the likelihood.

Finally, we reconstruct ancestral compositions of amino acid on alignments composed of buried residues only, we show the strongest correlation with the thermostability of the proteins. The reconstruction is performed on a anlignment of genes which are not strongly associated with one or another topology. The reconstruction with a model accounting for amino acid variation over time is given in the main text. Supplementary figure 9 displays the reconstruction with a standard evolutionary model that does not account for the variation.

Overall, the two methods of reconstruction are in strong aggreement due to the anchoring of compositions with modern species. Indeed, for the most recent ancestors attached to several modern species by short evolutionary time. In supplementary figure 9A, it appears that the model not taking into account amino acid variation (full dots) produces deep ancestors that are closer to the mean of the alignment (point of coordinates 0, 0), as it is expected because the equilibrium frequencies are calculated as the mean over the alignment. On the contrary, taking into account the variation allows reconstruction with greater flexibility, and all the ancestors (fainted dots) are shifted to the left of the graph, indicating that the mean compositions could be more similar to modern Alvinellidae in this model.
On supplementary figure 9B, we see that the projection onto axis expected to correlate with the protein stability [10] (supplementary table 10) produces very similar results in the two models, taking into account or ignoring past amino acid variations. The difference between the two models is more noticable for the oldest ancestors characterized by longer evolutionary times.
Overall, the model accounting for amino acid variation predicts ancestors with amino acid compositions closer to the extant Alvinellidae. This is especially noticeable for the LCA between Amphartidae+Alvinellidae and the deepest Terebelliformia ancestors, which could have had slightly more thermotolerant ancestors than the predictions of the model without amino acid variations (the sequences expected to be more thermostable should be in the top right corner of supplementary figure 10.B).

| Amino acid | Index 1 coefficient | Index 2 coefficient | Properties |
|---|---|---|---|
| A | -0,04 | 0,04 | Non polar, small |
| R | -0,12 | -0,24 | (+) charged, large |
| N | -0,25 | -0,07 | Polar, medium |
| D | -0,34 | 0,06 | (-) charged, medium |
| C | 0,11 | 0,04 | Non polar, small |
| Q | -0,15 | -0,17 | Polar, medium |
| E | -0,23 | -0,09 | (-) charged, medium |
| G | -0,26 | 0,15 | Non polar, small |
| H | -0,10 | -0,18 | (+) charged, large |
| I | 0,40 | 0,15 | Non polar, large |
| L | 0,33 | 0,05 | Non polar, large |
| K | -0,15 | -0,20 | (+) charged, medium |
| M | 0,21 | -0,06 | Non polar, large |
| F | 0,26 | -0,10 | Non polar, aromatic, large |
| P | -0,15 | 0,82 | Non polar, small |
| S | -0,18 | -0,03 | Polar, small |
| T | -0,01 | -0,00 | Polar, medium |
| W | 0,17 | -0,15 | Non polar, aromatic, large |
| Y | 0,17 | -0,19 | Polar, aromatic, large |
| V | 0,34 | 0,17 | Non polar, medium |

Supplementary figure 10 - Amino acid coefficients on the first two PCA axis identified in Tsuboyama et al. [10]. Positive coefficients correlating with stability are in green, negative in red.

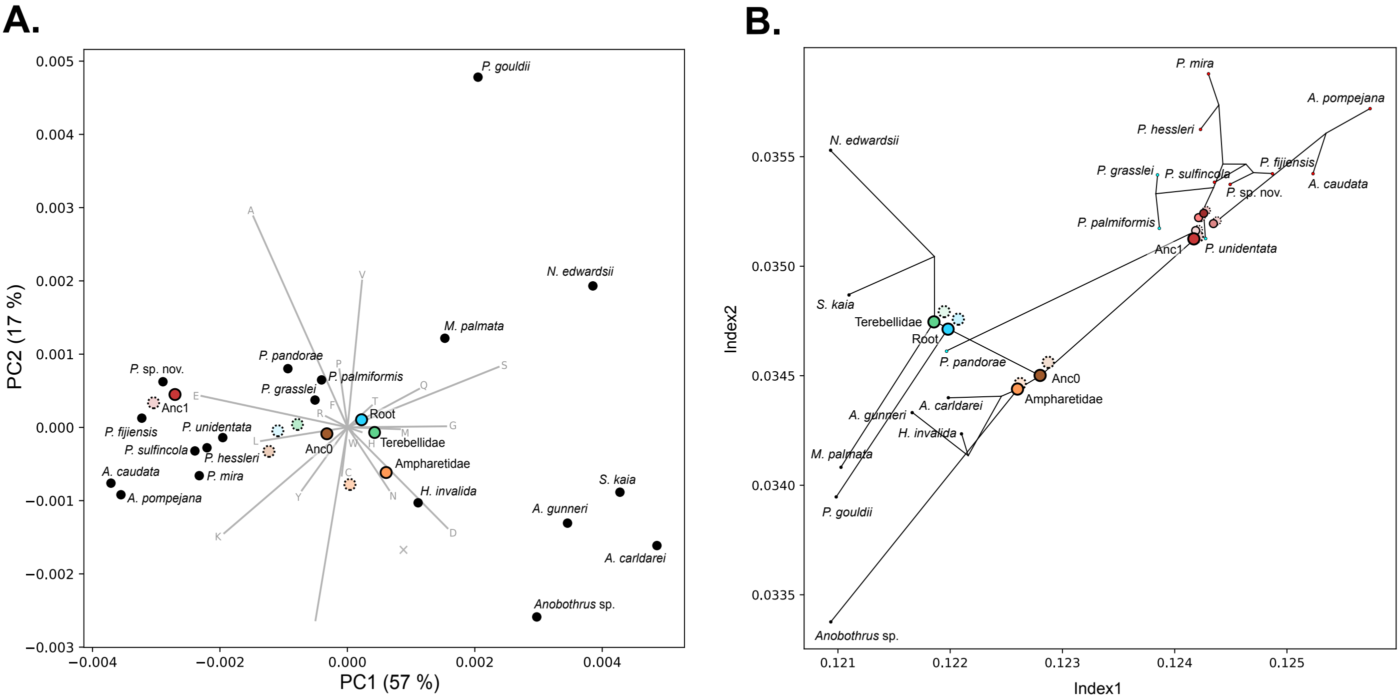


Supplementary figure 10 - Reconstruction of amino acid composition for buried residues, without taking into account the varying bias. Relative position of important ancestors are indicated in full (no variation in amino acid composition) or fainted/dotted (with variation in amino acid composition, as shown in the main article). A. PCA on the modern species. B. Projection on the axes of protein stability identified in Tsuboyama et al. [10].
